# Contribution of brain intrinsic branched-chain amino acid metabolism in a novel mouse model of maple syrup urine disease

**DOI:** 10.1101/2024.08.22.609255

**Authors:** Amanda C. Kuhs, Laura Ohl, Tegan Thurston, Jeet Singh, Sangeetha Bhuyan, Sarah Grandinette, Jing Xu, Sophie A Siemsgluess, Youseff Jakher, Rebecca C. Ahrens-Nicklas

## Abstract

Maple syrup urine disease (MSUD) results from loss of branched-chain ketoacid dehydrogenase (BCKDH) activity, the committed, rate-limiting step of branched-chain amino acid (BCAA) oxidation. Current treatments, including a low protein diet and liver transplantation, improve peripheral biochemistry and limit episodes of metabolic decompensation but do not fully prevent chronic neuropsychiatric symptoms. The mechanisms underlying chronic neurologic phenotypes remain poorly understood. Currently available MSUD mouse models do not survive long enough to evaluate chronic central nervous system (CNS) pathology. To investigate if loss of brain-intrinsic BCAA metabolism contributes to chronic neurologic disease, we developed a new brain-specific knockout mouse model of MSUD. First, we generated a mouse harboring a floxed *Dbt* allele (*Dbt*^flox/flox^). Then we crossed this line with Cre recombinase driver lines to induce loss of *Dbt* expression in 1) all developing CNS cell populations 2) neurons alone or 3) astrocytes alone. We found that brain-specific KO mice have elevations in BCAA levels in cortex that are exacerbated by a high protein diet. They also have secondary changes in amino acids that are important for neuronal function, including glutamine and glycine. These metabolic differences result in subtle functional deficits as measured by electroencephalogram and behavioral testing. Astrocyte and neuron-specific KO mice each also demonstrate mild biochemical features of MSUD in the cortex, suggesting that both cell populations may contribute to disease pathology. Collectively, these data suggest that therapies targeting the CNS directly, in addition to the periphery, may improve outcomes in MSUD.

## Introduction

Maple Syrup Urine Disease (MSUD) is a rare autosomal recessive genetic disorder that results from loss of branched-chain ketoacid dehydrogenase (BCKD) activity. BCKD catalyzes the committed step of branched-chain amino acid (BCAA) oxidation. MSUD is estimated to occur in 1:200,000 births^1–3^. In isolated populations the incidence is higher. For example, the Mennonite population of southeastern Pennsylvania has an incidence of approximately 1:360^4^. Without intervention MSUD causes encephalopathy, the progression of which can lead to coma and/or death^5,6^. Treatments for MSUD include dietary therapy and liver transplantation. Alongside the initiation of newborn screening, these interventions have improved the outcomes and lifespan of patients with MSUD^7–9^.

Liver transplantation improves peripheral biochemistry and largely eliminates episodes of metabolic decompensation in MSUD patients. However, longitudinal studies indicate that even this drastic intervention fails to halt the progressive neurologic, cognitive, and psychiatric decline that may reflect ongoing abnormal BCAA metabolism within the brain. Adolescents with MSUD are more likely to develop neuropsychological symptoms including depression, anxiety, and attention deficit disorder^6^. In a study of 37 classical MSUD patients (26 on dietary therapy and 11 status post liver transplantation), there was an 83% cumulative lifetime incidence of psychiatric conditions including depression, anxiety, inattention, and hyperactivity. Similarly, patients with MSUD had lower full-scale IQs as compared to controls^10^. A high burden of neuropsychiatric disease including deficits in attention and executive functioning has also been reported in several other treated MSUD cohorts^11,12^.

Investigations into the mechanisms of brain dysfunction in MSUD are ongoing. The prevailing hypothesis is that BCAA accumulation drives pathophysiology. Approximately 30-50% of the alpha-amino groups of glutamate and glutamine in the brain are derived from BCAAs^13,14^. During an episode of acute MSUD decompensation, serum and brain leucine levels increase and induce accumulation of branched-chain ketoacids. These ketoacids can be transaminated back to their parent amino acids, depleting the brain of glutamate, glutamine, aspartate, and alanine^14^. Thus, disruptions of BCAA metabolism likely fundamentally alter neurotransmission^15^. In addition, cerebral edema can occur.

The exact mechanisms underlying chronic neuropsychiatric symptoms in MSUD are still poorly understood, but sustained alterations in intrinsic CNS metabolism likely contribute to pathology. This hypothesis is supported by magnetic resonance spectroscopy (MRS) studies of well-controlled MSUD patients after liver transplantation which demonstrate persistent reductions in CNS glutamate, the major excitatory neurotransmitter of the brain^10^.

Two mouse models of MSUD have been developed^16^. The classic MSUD (cMSUD) knockout model has a deletion spanning exons 4 and 5 of *Dbt* which encodes the E2 subunit of the BCKDH complex. Knockout cMSUD mice die shortly after birth. The intermediate MSUD (iMSUD) model, engineered onto the background of the cMSUD model, has partial restoration of the E2 complex in the liver. This partial rescue of *Dbt* expression increases survival; however, most mice still die by early adulthood. Therefore, it is difficult to use these models to investigate mechanisms of chronic neurologic and psychiatric symptoms in patients with treated MSUD.

To address this unmet need, we have developed a new conditional knockout MSUD mouse model that harbors biallelic, floxed, *Dbt* alleles. When crossed to Cre recombinase driver lines, cell-type specific MSUD mouse models can be generated. Here, we utilize this novel model to investigate the mechanisms underlying chronic neurologic dysfunction in MSUD.

## Methods

### Mouse generation

All animal usage and experiments were approved by the Institutional Animal Care and Use Committee at the Children’s Hospital of Philadelphia (CHOP) in accordance with the US Public Health Service’s Policy on Humane Care and Use of Laboratory Animals.

Conditional knockout (*Dbt*^Flox/Flox^) mice^17^ were generated by the University of Pennsylvania CRISPR/Cas9 Mouse Targeting Core and CHOP Transgenic Core using CRIPSR/Cas9 (**Figure 1A and Supplementary Figure 1**). LoxP sites flanking exon 2 of *Dbt* were generated using the Alt-R™ CRISPR-Cas 9 genome editing system (Integrated DNA Technologies, Coralville, IA). The target sequences were as follows: 5′ - GTGAAGGTTATTGACAGAGTGGG; 3′ - CATTGTGGGAAACTATGGAGCGG. CRISPR guides were designed with the sequences as follows: 5′ - GTGAAGGTTATTGACAGAGT; 3′ - CATTGTGGGAAACTATGGAG. Trans-activating crRNA (tracrRNA, Integrated DNA Technologies, #1072533) was hybridized to crRNA to activate the Cas9 enzyme. After oocyte injection, founders were confirmed via Sanger sequencing of genomic DNA. Founders were backcrossed to C57BL/6J mice. Mice were then bred to homozygosity (*Dbt*^Flox/Flox^).

**Figure 1:**
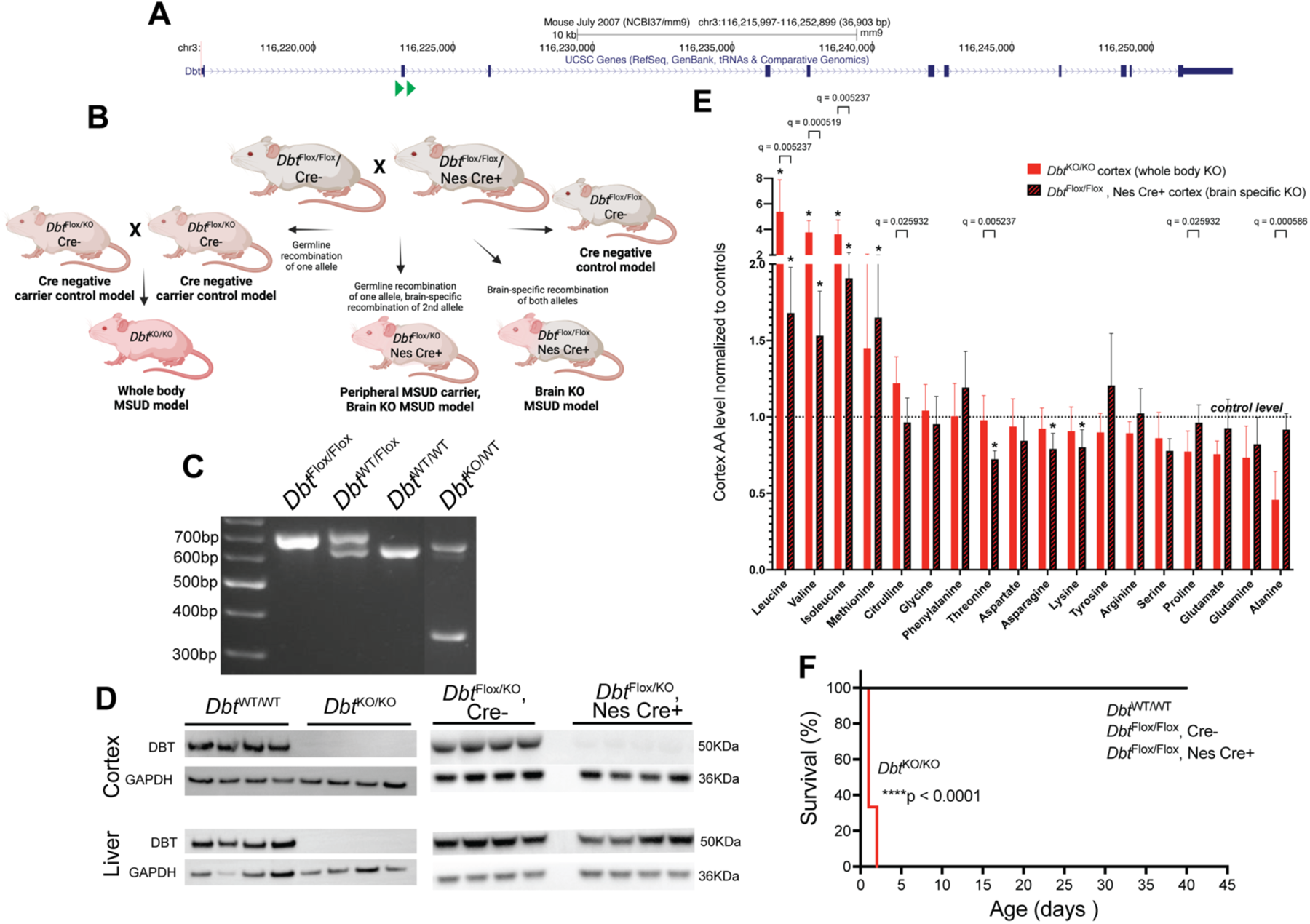
Brain-specific MSUD mice have elevated BCAA levels in cortex but normal survival. **(A)** LoxP sites flanking exon 2 of the *Dbt* gene were introduced to create *Dbt* floxed mice, see Supplementary Figure 1 for full sequence. **(B)** *Dbt*^Flox/Flox^ mice were crossed to mice harboring a Nestin-Cre allele to generate a brain-specific KO model of MSUD (*Dbt*^Flox/Flox^ , Nes Cre+). Due to high rates of germline recombination a substantial number of peripheral carrier, brain KO MSUD mice (*Dbt*^KO/Flox^ , Nes Cre+) were also generated. Finally whole body *Dbt* KO mice were generated by crossing heterozygous *Dbt*^KO/Flox^ mice. **(C)** PCR-based genotyping differentiates the *Dbt*^WT^ , *Dbt*^KO^ , and *Dbt*^Flox^ alleles. **(D)** DBT protein, as detected by western blotting) was absent in both the cortex and liver of whole body MSUD mice (*Dbt*^KO/KO^). However, DBT protein but is present in the liver but absent in the cortex of peripheral carrier, brain KO MSUD mice. **(E)** Amino acid levels in cortex were measured a p0-p1 in *Dbt*^KO/KO^ mice (red bars) and normalized to *Dbt*^WT/WT^ levels (control level indicated by dotted line). In addition, amino acid levels in cortex from p30 brain KO MSUD mice (*Dbt*^Flox/Flox^ , Nes Cre+, red and black striped bars) were measured and normalized to levels from age matched control (*Dbt*^Flox/Flox^ , Cre-) cortex (control level indicated by dotted line). Asterix indicate metabolites that differed from control values, as measured by multiple t-tests with multiple comparison analysis using the two-stage step-up method of Benjamini, Krieger and Yekutieli with a desired q of 5.00%. In addition, fold changes in whole body MSUD mice were compared to brain KO MSUD mice using multiple t-tests with multiple comparison analysis, and significant q values are shown. Data shown as mean with standard deviation. N=7 animals per genotype group. **(F)** Whole body MSUD mice (*Dbt*^KO/KO^ ) survive for a median of 1 day, which is significantly reduced as compared to brain KO MSUD mice and control animals (****p<0.0001, log-rank Mantel-Cox test). N= a minimum of 6 animals per genotype group.

To generate brain-specific *Dbt* knockout mice, *Dbt*^Flox/Flox^ mice were crossed with mice expressing Cre recombinase under control of the Nestin promoter (Nes-Cre, Jackson Laboratory, Bar Harbor, ME; B6.Cg-Tg(Nes-cre)1Kln/J, Stock No: 003771) (**Figure 1B**), which drives Cre expression throughout the developing central nervous system^18^. Nes-Cre mice also demonstrate high rates of germline recombination^19^, which facilitated generation of a full-body knockout mouse, *Dbt*^KO/KO^ (**Figure 1B**). For astrocyte specific knockout, *Dbt*^Flox/Flox^ mice were crossed to GFAP-Cre driver mice^20^ (Jackson Labs, B6.Cg-Tg(Gfap-cre)77.6Mvs/2J, Strain #024098), For neuron specific knockout, *Dbt*^Flox/Flox^ mice were crossed to Syn1-Cre driver mice^21^ (Jackson Labs, B6.Cg-Tg(Syn1-cre)671Jxm/J, Strain #003966). Unless noted, mice were maintained on a 17% protein, regular diet (LabDiet, #5015).

For studies of diet interventions, mice were transitioned to either a control diet with 20% of calories coming from protein diet (Envigo Teklad, Madison WI, #TD.91352) or a nutrient-matched high protein diet with 40% of calories coming from protein (40% of calories from protein, Envigo Teklad, Madison WI, #TD.90018). Animals were transitioned to these diets for 2 weeks prior to behavioral testing at p60 +/-7 days, followed by tissue harvest for metabolic testing at p70 +/-7 days. All diets were fed ad libitum.

### Genotyping

Genotyping was completed for the *Dbt* and Cre alleles as previously reported^17^ (See **Supplementary Methods**).

### Western blotting

Whole brain or liver tissue was briefly homogenized in RIPA buffer followed by BCA assay (ThermoFisher Scientific) to determine protein concentration. Equal amounts of protein from WT and KO conditions were loaded in the NuPAGE 4-12% Bis-Tris Protein Gels (ThermoFisher Scientific). Proteins were then transferred onto a nitrocellulose membrane (ThermoFisher Scientific) and blocked for 1 hour at room temperature (RT), followed by incubation in the primary antibody overnight at 4°C. After incubation with HRP-conjugated secondary antibodies, blots were developed with SuperSignal™ West Pico PLUS reagents (ThermoFisher Scientific) on a ChemiDoc Gel Imaging System BioRad, Hercules, CA). Blots were then analyzed in Image J software (NIH). The following antibodies and dilutions were used: 1:6,000 for DBT (Proteintech, #12451-1-AP), and 1:5,000 for GAPDH (Proteintech, #60004-1).

### Amino acid analysis from p0 cortex

For *Dbt* ^KO/KO^ amino acid analysis, offspring of *Dbt* ^WT/KO^ x *Dbt* ^WT/KO^ crosses underwent cryoanesthesia and decapitation. Cortices were collected and flash frozen in liquid nitrogen on postnatal day 1. Amino acid concentrations were determined by The CHOP Metabolomics Core with an Agilent 1260 Infinity HPLC system, utilizing pre-column derivatization with o-phthalaldehyde using previously published methods^22,23^.

### Metabolic analysis of adult cortex

For metabolomic analysis of cortex, p30 +/- 7 day or p70 +/- 7 day animals were anesthetized with isoflurane and underwent cervical dislocation followed by decapitation. Cortices were dissected and flash frozen in liquid nitrogen. Tissues were sent to the Penn Cardiovascular Institute Metabolomics Core. Wet frozen tissues were received on dry ice and stored at -80°C. The frozen tissues were lyophilized overnight.

For targeted metabolomics, dry tissue powders of each sample were homogenized in 50/50 0.3% formic acid/acetonitrile. An aliquot of each homogenate was prepared for amino acid and acylcarnitine analysis by LC/MS, according to validated, optimized protocols in previously published studies^17,24,25^. These protocols use cold conditions and solvents to arrest cellular metabolism and maximize the stability and extraction recovery of metabolites. Each class of metabolites was separated with a unique HPLC method to optimize their chromatographic resolution and sensitivity. Quantitation of metabolites in each assay module was achieved using multiple reaction monitoring of calibration solutions of metabolites and study samples spiked with matched isotopically-labelled internal standards (**Supplementary Table 1 and 2**) on an Agilent 1290 Infinity UHPLC/6495 triple quadrupole mass spectrometer. Raw data was processed using Mass Hunter quantitative analysis software (Agilent). Calibration curves (R^2^ = 0.99 or greater) were fitted with either a linear or a quadratic curve with a 1/X or 1/X^2^ weighting.

For untargeted metabolomics, frozen, powdered samples were weighed and homogenized in ice cold 80% methanol (homogenate density of 20 mg/mL) in a Precellys homogenizer at 4 °C. Two 100 µL aliquots of each homogenate (one each for reversed-phase C18 chromatography and HILIC chromatography) were extracted with 400 µL of ice cold methanol, vortexed, centrifuged at 18,000 x g for 5 minutes at 4 °C, and 400 µL of the supernatants were dried down under nitrogen at 45 °C in a 96-well plate. An aliquot of each brain homogenate was also combined to make pooled QCs for each chromatography mode and similarly prepared. Dried samples were reconstituted in 200 µL of HPLC solvents in the 96-well plate, and reversed-phase C18 chromatography was performed to retain and separate medium polarity to nonpolar metabolites on a Thermo Scientific Vanquish UHPLC. HILIC chromatography was performed to retain highly polar metabolites not retained by reversed-phase C18 chromatography. The Vanquish UHPLC was coupled to an Orbitrap ID-X mass spectrometer scanned from *m/z* 60-1000 at a resolution of 120,000. Compound Discoverer (Thermo Scientific) was used to process the LC/MS metabolomics data to identify metabolites and determine statistical differences between study groups.

Untargeted metabolomics data were analyzed using MetaboAnalyst v.5^26^ (https://genap.metaboanalyst.ca/home.xhtml). To evaluate if the metabolite peak patterns clustered by genotype groups, hierarchical clustering using Euclidian distance as a similarity measure and Ward’s linkage clustering algorithm were used to generate a heatmap and dendrogram. In addition, partial least squares discriminate analysis (PLS-DA) was completed. This supervised method uses multivariate regression techniques to extract information that can predict class membership. Two-dimensional PLS-DA plots were generated.

An input peak table containing m/z values and retention times sorted by p-values was created for functional analysis. To investigate pathway-level differences between *Dbt*^flox/flox^ Cre negative controls and brain-specific *Dbt* knockout mice, the MetaboAnalyst peaks-to-pathways module was used to analyze peaks from both C18 and HILIC experiments in both positive and negative ionization mode. This module performs a meta-analysis of mummichog and GSEA pathways. Specifically, the module uses the Fisher’s method for combining the mummichog and GSEA p-values. It takes the raw p-values per pathway to perform p-value combination. Analysis was completed using a Mummichog algorithm with a p-value cutoff of 1.0E-5 and the Mus musculus KEGG pathway library for annotation.

### Immunohistochemistry

Brain sectioning and immunohistochemistry were performed as previously described^27^ (also see **Supplementary Methods**).

### Electroencephalogram (EEG)

EEG recording electrodes were constructed and implanted as previously described^27,28^ into p60 mice (see **Supplementary Methods**).

### Behavioral assays

Mice were acclimated to the testing room in their home cages for 30 minutes before the start of any assay. When assays were performed on the same day, mice were allowed to reacclimate for at least 30 minutes prior to the start of the next assay.

*Open field testing:* The open field apparatus is a white rectangular box (60 cm x 50 cm x 26 cm), consisting of an outer region and an inner region. Mice were placed in the lower right corner of the open field apparatus and allowed to explore, undisturbed, for 15 minutes while an overhead video camera recorded their behavior. For analysis, the total distance traveled, mean speed, time and entries in the inside and outside regions were calculated using ANY-maze software (ANY-maze, Wood Dale, IL).

*Elevated zero testing:* The elevated zero apparatus is a circular raised platform (2 feet from the ground) with a diameter of 60 cm, consisting of two open arms opposed to each other and two high-walled closed arms opposite of each other. Mice were placed in the closed arm of the apparatus and allowed to explore, undisturbed, for 5 minutes. The mice were monitored and recorded using an overhead video camera. For analysis, total distance traveled, mean speed, time and entries in the open arms versus closed arms were calculated using ANY-maze software.

### Statistical analysis

Data are presented as mean ± SEM. Statistical analysis was performed using Prism software (GraphPad, San Diego, CA). The respective statistical test for each experiment is indicated in the figure legends. Data was tested for normality and nonparametric methods were used as appropriate. Significance is denoted in each figure. Both male and female mice were used for all experiments, and data were analyzed in a blinded fashion when possible.

## Results

### Generation and validation of a brain-specific MSUD mouse model

Mice harboring loxP sites flanking exon 2 of *Dbt* (*Dbt* ^Flox/Flox^ ) were successfully generated (**Figure 1A-C, Supplementary Figure 1**). These mice were then crossed to mice carrying the Nestin Cre allele (Nes Cre+) to generate a brain-specific (i.e., “brain KO”) model of MSUD (**Figure 1B**). Due to high rates of germline recombination in Nes Cre+ mice^19^, both whole body MSUD and peripheral carrier / brain KO MSUD models were also generated (**Figure 1B)**. Peripheral carrier / brain KO MSUD mice can serve as a model of partially treated MSUD such as occurs in patients after liver transplantation. In this case, peripheral, but not brain, BCAA metabolism is partially improved. Whole body KO MSUD mice lacked Dbt protein in both the brain and liver as detected by western blotting. Conversely, peripheral carrier / brain KO MSUD mice expressed Dbt protein in liver, but not cortex (**Figure 1D)**.

Targeted amino acid analysis of cortex from whole body KO MSUD mice (**Figure 1E, red bars)** demonstrated a 5.4, 3.8 and 3.6-fold increase in leucine, valine, and isoleucine, respectively, as compared to wild-type mice. Brain KO MSUD (*Dbt*^Flox/Flox^, Nes Cre+) mice (**Figure 1E, black / red bars)** had a more subtle increase in BCAAs in cortex with a 1.7, 1.5, and 1.9-fold increase in leucine, valine, and isoleucine, respectively, as compared to *Dbt*^Flox/Flox^, Cre-control cortex.

In brain, BCAAs serve as a major nitrogen donor in the synthesis of glutamate, glutamine and alanine^29^. There was a trend towards reduced levels of these key CNS amino acids in whole body MSUD mice as compared to wild-type cortex, consistent with previously-reported reductions in the blood of a classic MSUD mouse model^16^. However, it is important to note that there was significant animal-to-animal variability in cortical amino acids in our p0 cohort. Cortex from brain KO mice did not have a significant decrease in glutamate, glutamine or alanine as compared to *Dbt*^Flox/Flox^, Cre-control cortex.

Whole body KO MSUD mice died shortly after birth, with a median survival of 1 day, consistent with a previously reported classic MSUD mouse model. Brain KO and peripheral carrier / brain KO MSUD mice demonstrated normal survival (**Figure 1F)**.

### Metabolic abnormalities extend beyond the BCAA pathway in brain-specific MSUD mice

To further investigate metabolic differences in the brain KO MSUD mouse model, untargeted metabolomic analysis was completed via LC/MS analysis of cortical tissue. Metabolomic profiles from brain KO MSUD mice and control mice clustered by genotype by both partial least squares discriminate analysis and hierarchical clustering (**Figure 2A-B)**. A meta-analysis of mummichog and GSEA pathway analysis confirmed that BCAA amino acid degradation was the most disrupted pathway in the cortex of brain KO MSUD mice, validating our model. However, additional pathways were disrupted, including aminoacyl tRNA biosynthesis and glycerophospholipid metabolism.

**Figure 2:**
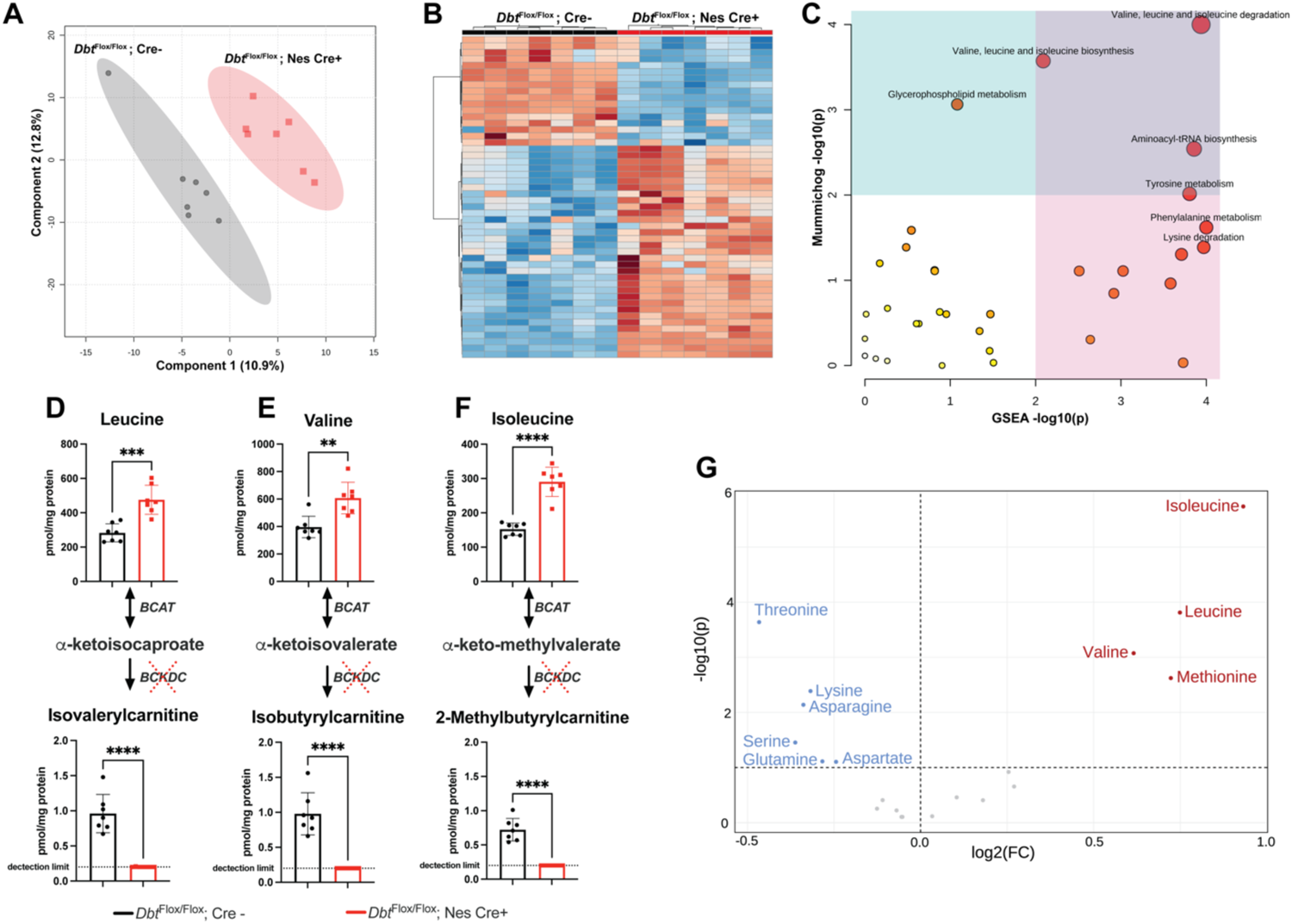
Brain-specific MSUD mice have abnormal global and targeted metabolomic profiles in adolescence. **(A)** Untargeted LC/MS metabolomic analysis was completed in p30 cortex from control (*Dbt*^Flox/Flox^, Cre-, indicated in black) and brain KO MSUD mice (*Dbt*^Flox/Flox^ , Nes Cre+, indicated in red). Metabolic profiles clustered by genotype when analyzed through partial least squares discriminate analysis. **(B)** In addition, profiles clustered by genotype when analyzed by hierarchical clustering using Euclidian distance as a similarity measure and Ward’s linkage clustering algorithm. These data were used to generate a metabolite heatmap and dendrogram. **(C)** A meta-analysis of mummichog and GSEA pathway analysis reveals that BCAA degradation is the most perturbed metabolic pathway in brain KO MSUD mice; however, additional pathways are also perturbed. The size and color of the circles correspond to their transformed combined p-values. Large, red circles are considered the most altered pathways as compared to controls. The blue and pink areas highlight the significant pathways based on either GSEA (pink) or mummichog (blue), and the purple area highlights significant pathways identified by both algorithms. **(D-F)** Targeted metabolomic data confirms an increase concentration in all BCAAs and decrease in downstream acylcarnitine species in p30 brain KO MSUD mice (red) as compared to controls (black). Data shown as mean +/-standard error, and groups were compared via student’s t-test with **p<0.01, ***p<0.001, ****p<0.0001. **(G)** Volcano plot demonstrating perturbations of additional amino acids beyond BCAA on targeted analysis.

To validate the untargeted metabolomic analysis, targeted amino acid and acylcarnitine profile analysis of p30 cortex was completed. BCAA levels were elevated in the cortex of brain KO MSUD mice (**Figure 1** and **2 D-G**). In addition, there were significant reductions in downstream acylcarnitine species including isovalerylcarnitine, isobutyrylcarnitine, and 2-methyl-butyrylcarnitine. There were also secondary changes in additional amino acids (**Figure 2G**); however, no changes were identified in expression level of the key amino acid transporters, *Slc7a5* and *Slc38a2* (**Supplementary Figure 2**).

### High protein diet exacerbates metabolic abnormalities in the cortex of brain-specific MSUD mice

Patients with MSUD who undergo liver transplantation or are treated with strict dietary control have only partial correction of peripheral metabolism. Furthermore, brain-intrinsic BCAA metabolism is not corrected by these therapies. To model this condition, we generated peripheral carrier, brain KO MSUD mice (*Dbt*^KO/Flox^ , Nes Cre+). As compared to wildtype control (*Dbt*^Flox/Flox^, Cre-), and MSUD carrier (*Dbt*^KO/Flox^, Cre-) mice, peripheral carrier, brain KO MSUD mice had increased BCAA levels in cortex and decreased downstream acylcarnitine species on targeted metabolic analysis (**Figure 3A-C)**, with a 1.7, 1.4, and 1.8-fold increase in leucine, valine, and isoleucine, respectively at p70. These increases were further exacerbated when peripheral carrier, brain KO MSUD mice were placed on a 40% protein diet with a 2, 1.8, and 2.3-fold increase in leucine, valine, and isoleucine, respectively, as compared to *Dbt*^Flox/Flox^, Cre-control mice on a regular (20%) protein diet (**Figure 3A-C)**. Peripheral carrier, brain KO MSUD mice had additional amino acid abnormalities in cortex as compared to carrier (*Dbt*^KO/Flox^, Cre-) controls, including reduced glycine, asparagine, and glutamine (**Figure 3D, Supplementary Figures 3-4)**. A high protein diet exacerbated other amino acid changes, such as lowering alanine and arginine levels.

**Figure 3:**
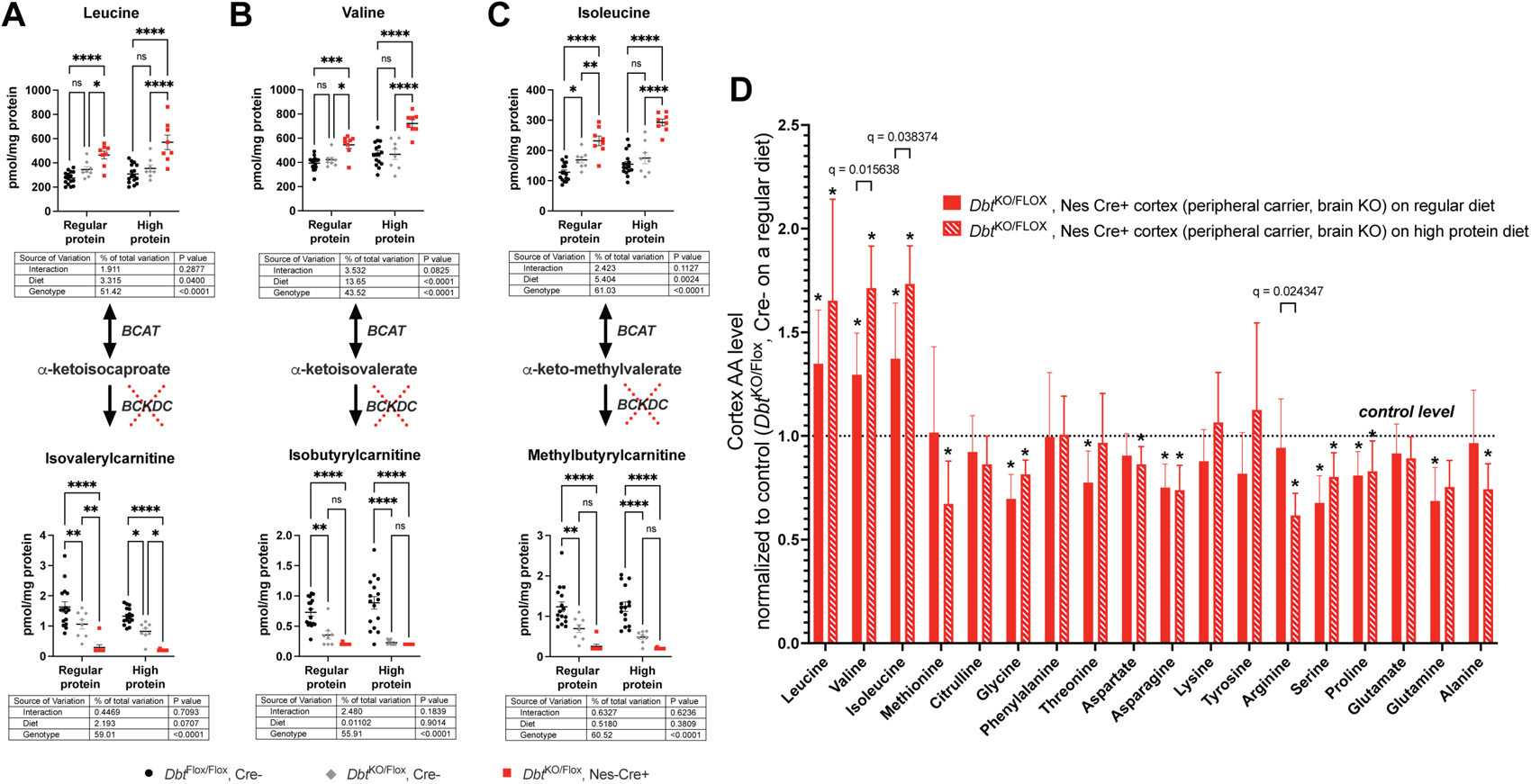
A high protein diet exacerbates metabolic abnormalities in the cortex of adult peripheral carrier, brain KO MSUD mice. P70 peripheral carrier, brain KO mice were investigated as a model of partially-treated MSUD (such as occurs after liver transplantation). **(A-C)** Targeted amino acid and acylcarnitine levels were compared between control (*Dbt*^Flox/Flox^, Cre-, black), peripheral carrier (*Dbt*^Flox/KO^ , grey), and peripheral carrier, brain KO MSUD mice (*Dbt*^Flox/KO^ , Cre+, red) on regular (20%) and high (40%) protein diet. Carrier mice showed a mild increase in isoleucine levels in cortex and a reciprocal decrease in all downstream acylcarnitine species as compared to controls. Peripheral carrier, brain KO mice showed a robust increase in BCAA levels and decrease in downstream acylcarnitine levels that was worsened on a high protein diet. Data shown as mean +/-standard error, and groups were compared via 2-way ANOVA followed by Sidak’s multiple comparison testing with *p<0.05, **p<0.01, ***p<0.001, ****p<0.0001. **(D)** Amino acid levels in cortex of peripheral carrier, brain KO MSUD mice on a regular (20%) protein diet (red, solid bars) were compared to peripheral carrier, brain KO MSUD mice on a high (40%) protein diet (red, striped bars). Each group was normalized to *Dbt*^KO/Flox^, Cre-control mice, indicated by the dotted line. Asterix indicate metabolites that differed from control values, as measured by multiple t-tests with multiple comparison analysis using the two-stage step-up method of Benjamini, Krieger and Yekutieli with a desired q of 5.00% (**See also Supplementary Figures 3-4**). In addition, fold changes in each dietary group were using multiple t-tests with multiple comparison analysis, and significant q values are shown. Data shown as mean +/-standard deviation. N=8 animals per genotype group.

Despite these metabolic abnormalities, peripheral carrier, brain KO MSUD mice had no significant differences in brain size or morphology, including similar cortical thickness and neuronal number **(Supplementary Figure 5)**. To evaluate for more subtle structural differences that could contribute to MSUD-related neurologic dysfunction, we quantified the density and size of dendritic protrusions in the CA1 region of the hippocampus, a region essential for learning and memory **(Supplementary Figure 6)**. There were no differences in spine density or length between carrier mice and peripheral carrier, brain KO MSUD mice.

### Neuron and astrocyte-specific MSUD models have diet-dependent differences in brain metabolites

To elucidate the contribution of different CNS cell populations to neurological abnormalities in MSUD, *Dbt*^Flox/Flox^ mice were crossed to mice harboring a Synapsin1-Cre (Syn1 Cre+) allele to generate a “neuron KO” MSUD mouse (*Dbt*^Flox/Flox^ , Syn1 Cre+). Additionally, mice were crossed to a Gfap-Cre (Gfap Cre+) driver line to generate an “astrocyte KO” MSUD mouse (*Dbt*^Flox/Flox^ , Gfap Cre+) (**Figure 4A)**. Loss of Dbt protein expression in the relevant cell type was confirmed via immunohistochemistry. Dbt was not expressed in astroctyes in the astrocyte KO model (**Figure 4B**) and was not expressed in neurons in the neuron KO model (**Figure 4C**).

**Figure 4:**
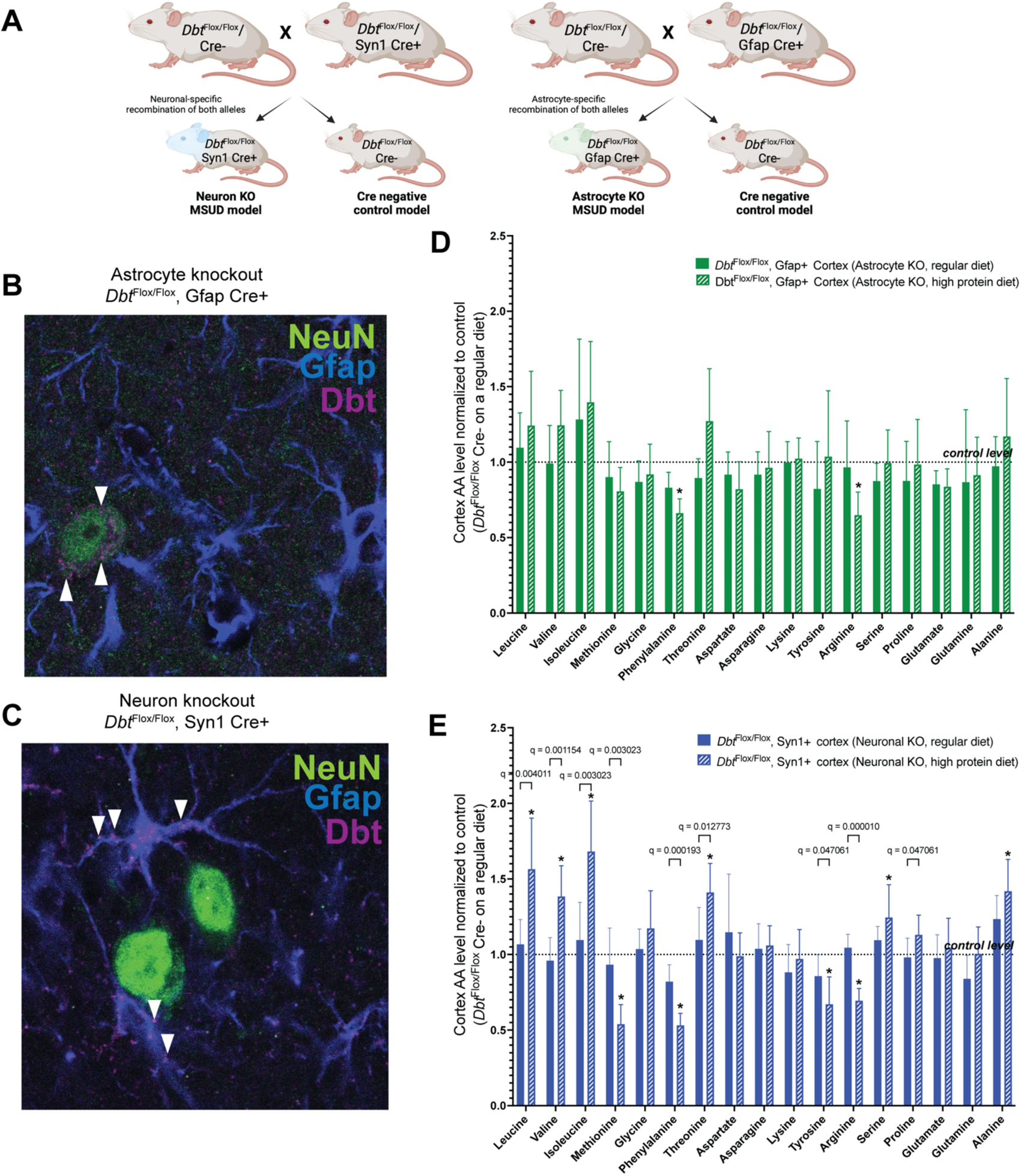
Successful generation of neuron-specific and astrocyte-specific KO MSUD mice. **(A)** *Dbt*^Flox/Flox^ mice were crossed to mice harboring a Synapsin1-Cre (Syn1 Cre+) allele to generate a neuron specific KO MSUD mouse (*Dbt*^Flox/Flox^ , Syn1 Cre+, indicated in blue). In parallel, *Dbt*^Flox/Flox^ mice were crossed to mice harboring a Gfap-Cre (Gfap Cre+) allele to generate an astrocyte specific KO MSUD mouse (*Dbt*^Flox/Flox^ , Gfap Cre+, indicated in green). **(B)** Representative immunofluorescence images of astrocyte KO MSUD cortex shows colocalization of Dbt (purple puncta, Dbt staining) with neurons (green, NeuN staining), as indicated by white arrow heads. While, Dbt positive puncta are absent from astrocytes (blue, Gfap staining). **(C)** Neuron KO MSUD cortex shows the reciprocal staining pattern, with colocalization of Dbt puncta (purple) with astrocytes (blue), as indicated by white arrow heads, but not neurons. **(D)** Amino acid levels in cortex of p70 astrocyte KO MSUD mice on a regular (20%) protein diet (green, solid bars) were compared to astrocyte KO MSUD mice on a high (40%) protein diet (green, dashed bars). **(E)** Amino acid levels in cortex of p70 neuron KO MSUD mice on a regular (20%) protein diet (blue, solid bars) were compared to neuron KO MSUD mice on a high (40%) protein diet (blue, dashed bars). For D-E, each group was normalized to *Dbt*^Flox/Flox^, Cre-control mice, indicated by the dotted line. Asterix indicate metabolites that differed from control values, as measured by multiple t-tests with multiple comparison analysis using the two-stage step-up method of Benjamini, Krieger and Yekutieli with a desired q of 5.00% (**See also Supplementary Figures 7-10**). In addition, fold changes in each dietary group were using multiple t-tests with multiple comparison analysis, and significant q values are shown. Data shown as mean +/-standard deviation. N=8 animals per genotype group.

Targeted amino acid analysis from cortex of astrocyte KO MSUD mice (**Figure 4D, Supplementary Figures 7-8)** revealed only a trend towards increased branched chain amino acid levels when mice were placed on a high (40%) protein diet. Of note, metabolites were measured from cortical homogenates which contain neurons, astrocytes, and a variety of additional CNS cell types. Cortex from neuron KO mice demonstrated more profound amino acid abnormalities as compared to astrocyte KO cortex, including elevated BCAA levels when challenged with a high protein diet (**Figure 4E, Supplementary Figures 9-10**). Interestingly, only neuron-specific KO mice recapitulated the low glycine levels that were seen in the peripheral carrier, brain KO mice. Both neuron and astrocyte KO MSUD mice recapitulated the low arginine levels while on high-protein diet that was seen in peripheral carrier, brain KO mice.

Untargeted metabolomic analysis was completed via LC/MS analysis of cortex. Metabolomic profiles from brain KO MSUD mice overlapped with astrocyte KO MSUD mice but differed from neuron KO MSUD mice by partial least squares discriminate analysis (**Figure 5A)**. Heatmaps from the 4 genotype groups segregated using hierarchical clustering (**Figure 5B)**. Follow-up targeted metabolic analysis confirmed only trend or minor increases in BCAA elevations and decreases in downstream acylcarnitine species in either astrocyte or neuron KO MSUD mice (**Figure 5C-E**). However, these subtle changes worsen when mice are placed on a high protein diet.

**Figure 5:**
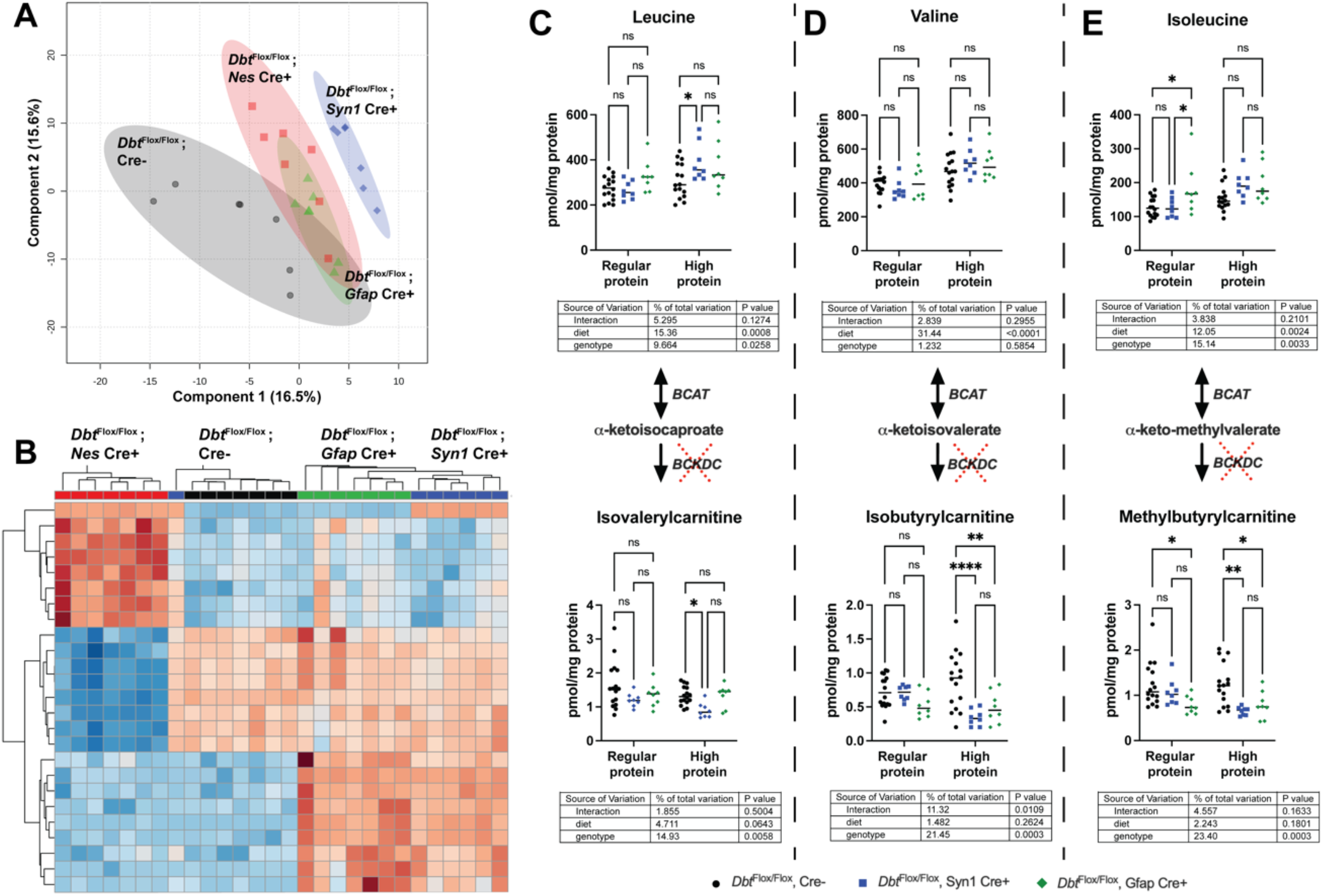
Neuron and astrocyte specific KO MSUD mice have abnormalities on global and targeted metabolic analysis that are exacerbated by a high-protein diet. **(A)** Untargeted LC/MS metabolomic analysis was completed in p30 cortex from control (*Dbt*^Flox/Flox^, Cre-, indicated in black), brain KO MSUD mice (*Dbt*^Flox/Flox^ , Nes Cre+, indicated in red), astrocyte KO MSUD mice (*Dbt*^Flox/Flox^ , Gfap Cre+, indicated in green), and neuron KO MSUD mice (*Dbt*^Flox/Flox^ , Syn1 Cre+, indicated in blue). Metabolic profiles clustered by genotype when analyzed through partial least squares discriminate analysis. Control and brain KO MSUD mice untargeted metabolite data (from Figure 2A-B) were reanalyzed to compare with neuron and astrocyte KO MSUD data here. **(B)** In addition, profiles clustered by genotype when analyzed by hierarchical clustering using Euclidian distance as a similarity measure and Ward’s linkage clustering algorithm. These data were used to generate a metabolite heatmap and dendrogram. **(C-E)** Targeted amino acid and downstream acylcarnitine levels were compared in cortex between all groups from p70 control (black), neuron KO MSUD (blue), and astrocyte KO MSUD (green) mice on regular and high protein diet. Data shown are mean +/-standard error, and groups were compared via 2-way ANOVA followed by Sidak’s multiple comparison testing with *p<0.05, **p<0.01, ***p<0.001, ****p<0.0001.

### Brain specific MSUD mice have hippocampal EEG and behavioral abnormalities

To evaluate the functional consequences of loss of *Dbt* expression in the brain, we completed longitudinal electroencephalogram (EEG) recordings from the CA1 region of the hippocampus of brain KO, neuron KO and astrocyte KO mice. We selected to focus on analysis of hippocampal recordings as it is a region implicated in neurocognitive and neuropsychiatric disorders. Brain KO MSUD mice had abnormal background frequency composition on EEG recordings (**Figure 6A-B)**, with decreased raw power in the slow, delta (0.1-4Hz) frequency range. Neuron (**Figure 6C)** and astrocyte (**Figure 6D)** KO MSUD mice did not show the same decrease in delta power. Brain KO MSUD mice also had an increased rate of epileptiform spikes in the hippocampus (**Figure 6E)**.

**Figure 6:**
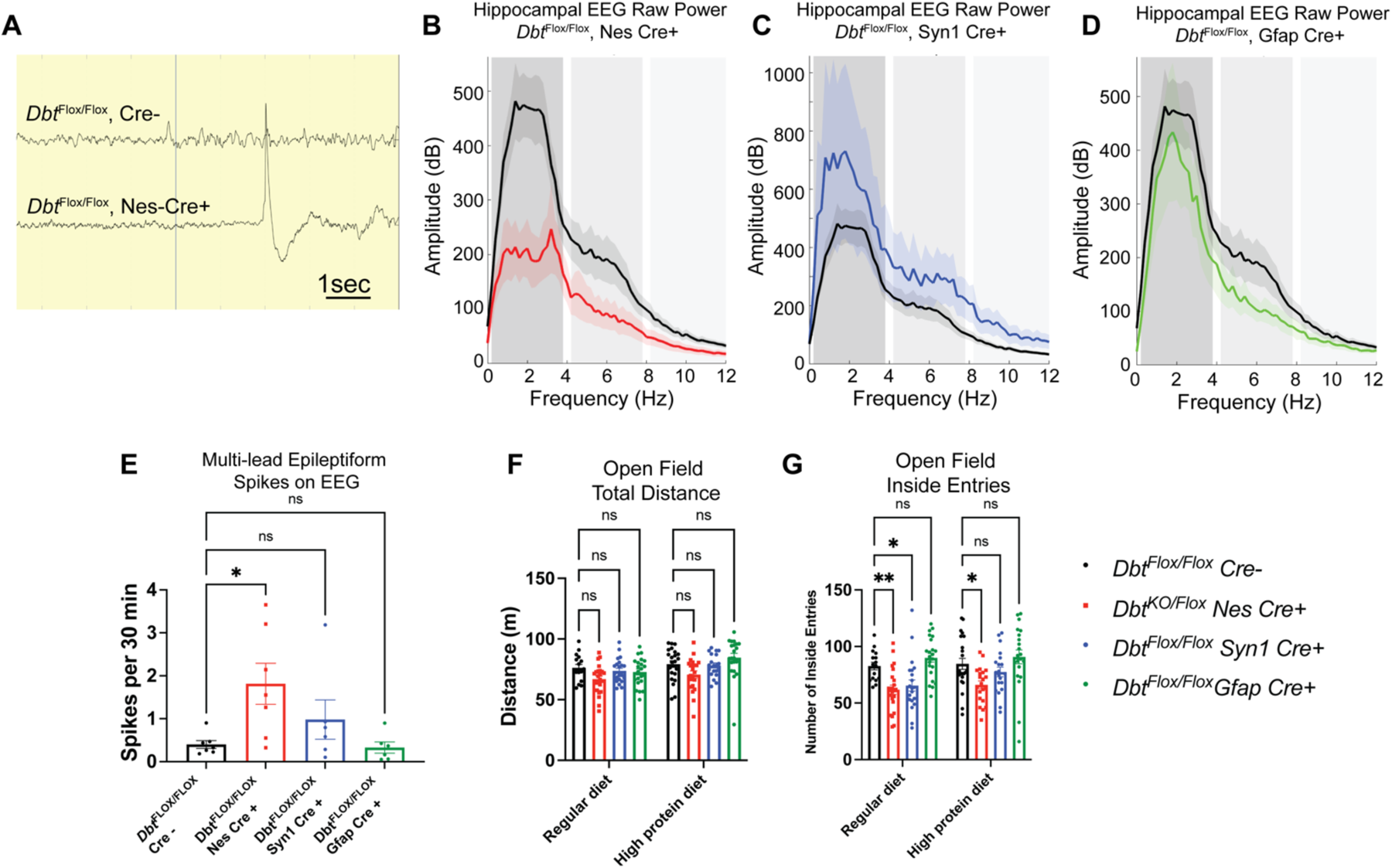
Brain KO MSUD mice have background abnormalities and increased epileptiform spikes on hippocampal EEG recordings and a subtle anxiety phenotype on behavioral testing. Longitudinal EEG analysis was completed on p60 brain KO, neuronal KO, and astrocyte KO MSUD mice. **(A)** Representative EEG traces recorded from the CA1 region of the hippocampus of show decreased slow delta power and an epileptiform spike in brain KO MSUD mice (*Dbt*^Flox/Flox^, Nes Cre+) as compared to controls (*Dbt*^Flox/Flox^, Cre-) **(B)** Fast Fourier transform analysis of 12 hours of EEG from brain KO MSUD mice (red) reveals a decrease in raw power in the slow delta band (0.1-4Hz) as compared to controls (black). Data shown as mean +/-standard deviation. **(C)** Neuron KO MSUD mice (blue) show a trend toward increased slow delta power, but the standard deviations of the two conditions overlap. **(D)** Astrocyte KO MSUD mice (green) do not differ from control mice in their raw power spectrum. **(E)** The number of multi-lead EEG spikes were quantified from 48 continuous thirty-minute recording periods (24 hours) per animal in control (black), brain KO (red), neuron KO (blue) and astrocyte KO (green) MSUD mice. Brain KO MSUD had an increased rate of epileptiform spikes as compared to the other genotypes. Groups were compared via nonparametric Kruskall-Wallis test followed by multiple comparison analysis with uncorrected Dunn’s test, *p<0.05. For EEG studies, N=6-7 animals per genotype. **(F)** None of the groups differed in their total distance traveled on open field testing on a regular or high protein diet. **(G)** Brain KO and neuron KO MSUD mice had fewer entries into the inside region of the open field chamber, suggestive of a mild anxiety phenotype, as compared to control or astrocyte KO MSUD mice. Groups were compared via 2-way ANOVA followed by Sidak’s multiple comparison testing with *p<0.05, **p<0.01. N= minimum of 16 mice per genotype group, balanced for sex.

Finally, mice were assessed for behavioral phenotypes with elevated zero and open field behavioral assays. In preliminary studies, we found that the Cre allele itself could influence behavioral phenotypes, consistent with previous reports of behavioral phenotypes in Cre driver lines^30^. We specifically found Cre-induced differences on the elevated zero assay, but only minimal effects on open field analysis **(Supplementary Figure 11)**; therefore, we proceeded with open field testing as a measure of motor activity and anxiety in our MSUD models. None of the groups differed in their total distance traveled on open field testing (**Figure 6F)**. However, brain and neuron KO MSUD mice demonstrated a mild anxiety phenotype with decreased entries into the inside zone on open field testing (**Figure 6G**).

## Discussion

Patients with chronic, treated MSUD have a high burden of neurologic and psychiatric disease^8,10,11^. This has been demonstrated even after implementation of newborn screening for MSUD. While strict peripheral metabolic control through either liver transplantation or dietary therapy clearly improves neurologic outcomes, neuropsychiatric symptoms still occur. The mechanisms underlying chronic neurologic dysfunction in MSUD remains poorly understood.

We aimed to develop a mouse model of MSUD that would facilitate investigations of chronic neurologic disease. Previously-developed MSUD models recapitulate classic phenotypic and biochemical features of disease, but do not survive long enough to study chronic disease^16^. Therefore, we generated a conditional *Dbt* knockout mouse model, which can be crossed to Cre recombinase lines to facilitate loss of BCKDH complex activity in specific cell types or specific developmental timepoints.

To evaluate the consequence of loss of BCKDH activity in all cell types in the brain, we first crossed our conditional *Dbt* model to a Nestin Cre driver line (i.e., brain KO MSUD mice). Unfortunately, the Nestin Cre mouse exhibits a high rate of germline recombination, in addition to the intended recombination of the floxed allele in CNS cells^19^. Therefore, one limitation of this work is that we had a limited population of pure brain KO mice for analysis and could not conduct behavioral testing in this cohort.

However, we were able to generate a sufficient number of mice for targeted and untargeted metabolomic analysis, and longitudinal EEG recordings. Brain KO MSUD mice demonstrated normal survival and moderate (1.5-2 fold) increases in brain BCAA levels. In addition, they had mild decreases in a number of additional CNS amino acids with a trend towards reduced glutamate and glutamine levels. BCAAs serve as the major nitrogen donor for the synthesis of glutamate and glutamine^14,29^ in the CNS. Both are reduced in the brains of MSUD patients. Glutamine serves as the precursor to glutamate, the major excitatory neurotransmitter in brain. Abnormalities in both amino acids are implicated in a wide range of neurologic and psychiatric disorders^31–33^. In particular, there is emerging literature that alterations in glutamine metabolism occur in many neurodegenerative conditions, especially those with associated mitochondrial dysfunction^34^.

On untargeted metabolomic analysis, brain KO MSUD mice had secondary abnormalities in additional metabolic pathways that are key to neurologic function including glycerophospholipid metabolism and the metabolism of other amino acids. This includes a reduction in brain threonine, which has been previously reported in the intermediate, iMSUD, mouse model^35^. Interestingly, the iMSUD model is reported to have elevated lysine levels in brain, while the brain KO MSUD mouse had reduced lysine levels. Collectively, these data highlight that loss of intrinsic CNS BCAA metabolism can induce attenuated patterns of metabolites associated with classic MSUD and other neurologic disorders.

These metabolic abnormalities were sufficient to induce abnormal network-level electrical activity in the hippocampus on EEG monitoring. Brain KO MSUD mice had increased interictal spikes and decreased in raw power in the hippocampus. However, it is important to note, only subtle EEG changes were noted and no overt seizures were detected. Additional studies are needed to further characterize functional deficits brain KO MSUD mice.

We also sought to investigate the real-world condition of patients with partial restoration of BCAA metabolism in the periphery, such as occurs after liver transplantation. To model this condition, we generated peripheral MSUD carrier, brain KO MSUD mice. These mice also demonstrated increased BCAA levels in brain with a reduction in downstream acylcarnitine species. They had decreased glutamine levels with a trend towards decreased glutamate and aspartate levels. Interestingly these mice also demonstrated reduced glycine levels in brain, which have not been reported in previous MSUD mouse models. As glycine acts as an inhibitory neurotransmitter in the CNS, this change may also contribute to the pathophysiology of chronic neurologic symptoms in MSUD. When challenged with a high-protein diet, additional amino acid abnormalities were noted. Specifically, methionine, arginine, and alanine levels were reduced in the brain of peripheral carrier, brain KO mice as compared to carrier control mice on a regular diet. These data suggest that a high protein diet can worsen metabolic abnormalities in the brain of MSUD mice, even when peripheral Dbt levels are at carrier levels.

The glutamate-glutamine cycle is a tightly coupled series of reactions in neurons and astrocytes, that regulates glutamate levels in brain to prevent toxic build-ups of glutamate after it is released at excitatory synapses^36,37^. In this model, glutamate is released from neurons and quickly taken up by neighboring astrocytes. Astrocytes can uniquely convert glutamate to glutamine via the glutamate synthase enzyme. Glutamine can then be transported back into neurons for conversion back to glutamate to replenish presynaptic stores of this key excitatory neurotransmitter. As there is no major uptake mechanism for glutamate and glutamine in brain, and loss occurs overtime, synthesis of these key amino acids must occur in the CNS. BCAAs are a major source of nitrogen groups for the synthesis of glutamate and glutamine in brain^13,14,29^.

By crossing our floxed *Dbt* mouse to either neuron or astrocyte-specific Cre driver lines, we could evaluate the how loss of BCAA metabolism in either cell population altered neurochemistry. These results may help guide therapy development, as not all modalities will target all cell types. For example, AAV9, a vector widely used in gene therapy clinical trials, predominately transduces glia, but not neurons in nonhuman primates^38^. We observed that untargeted metabolomic profiles from cortex of both neuron and astrocyte specific KO mice differed from control animals. Each model showed a mild increase in BCAAs levels and a reciprocal decrease in downstream acylcarnitine species in brain when challenged with a high protein diet. Neuron KO mice had a more profound amino acid differences than astrocyte KO mice, especially when challenged with a high protein diet. Similarly, neuron KO mice had a detectable, albeit subtle, anxiety phenotype, while astrocyte KO mice did not.

In summary, disruption of BCAA metabolism in brain alters neurochemistry in a mouse model of chronic MSUD, even when peripheral BCAA metabolism remains intact. Key alterations include increased BCAAs levels and changes in glutamine levels. These changes are exacerbated by a high protein diet or when the mouse is a peripheral carrier for MSUD. Collectively, these changes had only subtle functional consequences on EEG and behavioral assays. In addition, there was no difference in survival in our brain specific KO mice. The mild functional phenotypes observed in brain KO mice, suggests that abnormalities in peripheral metabolism (as opposed to brain intrinsic BCAA metabolism) may be the primary driver of overt chronic neurological symptoms in MSUD. Future studies are needed to evaluate for the presence of additional neurocognitive or other behavioral deficits in brain KO models of MSUD.

Prior studies and the normal survival observed in our model suggest that therapies which correct peripheral BCAA metabolism will have the most significant impact on MSUD. However, our results raise the possibility that the addition of a CNS-targeting strategy will provide an additional benefit in this population.

## Author contributions

Project conceptualization and design: RAN, LO, ACK, JS, TT, SB, JX, SAS, YJ

Data acquisition: LO, ACK, JS, TT, SB, JX, SAS, YJ, SG

Data analysis: RAN, LO, ACK, JS, TT, SB, JX, SAS, YJ, SG

Funding acquisition: RAN

Manuscript drafting: RAN, ACK, TT, JS, JX

Critical review of manuscript: RAN, LO, ACK, JS, TT, SB, JX, SAS, YJ

## Funding

This work was supported by The Children’s Hospital of Philadelphia Research Institute and the MSUD Family Support Group.

## Data availability

Data generated and analyzed from this study are included in this article and the supplementary information. Data supporting these studies findings are available from the corresponding author with reasonable request.

## Competing interest statement

RAN is an advisor to LatusBio and AskBio on projects unrelated to this manuscript. The remaining authors have no competing interests.

## Acknowledgements

We would like to thank the University. of Pennsylvania Cardiovascular Institute Metabolomic Core, the University of Pennsylvania CRISPR/Cas9 Mouse Targeting Core, the Children’s Hospital of Philadelphia Transgenic Core, and the Children’s Hospital of Philadelphia Metabolic Core.

## Supplementary Methods

### Genotyping

Genotyping was completed via PCR. Top Taq Master Mix Kit (QIAGEN, #200403) or KAPA2G Fast HotStart ReadyMix (Roche, Switzerland, #KK 5609) was used for PCR reactions. Mouse tail clips or tissues were isolated using DNeasy Blood & Tissue Kits (QIAGEN, Hilden, Germany, #69509). *Dbt* primers were designed to flank the 5’ and 3’ loxp sites to distinguish *Dbt*^Flox^ , *Dbt*^KO^ , and *Dbt*^WT^ alleles. The sequences for the forward and reverse primer are 5’-ACCGGAGCATCAGCCTAAA, and 3’-TGTGCACAAGGACATACAGG. The product sizes are 621 bp for the wild-type allele, 311 bp for the knockout allele, and 689 bp for the floxed allele. Primers to detect the Cre allele were used as reported by the Jackson Laboratory (Jackson Labs #003771, B6.Cg-Tg(Nes-cre)1Kln/J).

### Immunohistochemistry

Mice were anesthetized with isoflurane and transcardially perfused with saline and 4% paraformaldehyde (PFA) in cold 1X phosphate-buffered saline (PBS). Brains were collected and postfixed overnight at 4°C in 4% PFA/PBS. Brains were cryoprotected in 30% sucrose buffer for at least 24 hours then embedded in OCT. 20 μm coronal sections were cut using a Leica CM3050 S Cryostat and collected on slides which were stored at -80°C until use.

For immunostaining glass slides with sections were first dried at 37°C for 15 min and washed with 1X PBS for 15 min at room temperature (RT). Sections were then permeabilize with 0.1% TritonX-100 in PBS for 7 min, washed 3 times with 1X PBS at RT for 5 minutes each, and then blocked in 10% normal goat serum (NGS)/0.1% Triton-100X/1X PBS for one hour at RT. Sections were then incubated at 4°C overnight in primary antibody (anti-NeuN mouse (Millipore Sigma MAB377, 1:100), anti-GFAP chicken (Thermo Scientific PA110004, 1:500) or anti-DBT rabbit (Proteintech 12451-1-AP, 1:500)).

For immunofluorescence, sections were then incubated in secondary antibody for 30 min at room temperature (anti-mouse Alexa Fluor 488 (Molecular probes, Cell Signaling Technology (CST) 4408S, 1:500), anti-rabbit Alexa Fluor 594 (Molecular probes, CST 8889S, 1:500) and anti-chicken 680 (Invitrogen A32934, 1:500).

Confocal Images were acquired on SP8 confocal microscope using the LAS X software (Leica). Each image stack was acquired with Pixel display resolution of 1024x1024, 8-bit dynamic range, 63X objective with, numerical aperture (NA) 1.40 and 3x zoom. White light laser was used to obtain following laser line: 488 nm, 594 nm and 647nm. XY pixel size was ∼141 nm and focal plane interval was ∼300 nm. Image J (NIH) was used for the image processing.

For DAB detection of the NeuN primary antibody, slide were incubated with a biotinylated goat anti-mouse secondary antibody (Jackson ImmunoResearch, 115-065-062, 1:500) at RT for 30 minutes, followed by an avidin-biotin complex (Vector Laboratories, PK-6100) at RT for 1 hour, and finally a DAB solution ((Vector Laboratories, SK-4105) for 4 minutes, followed by PBS washes. Slides were washed with PBS as described above in between each of the previous steps. Slides were then washed two times with distilled water for two minutes. 0.1% cresyl violet in acetate buffer was added to the slides for 3 minutes, followed by a quick acid wash in ethanol/hydrochloric acid. Slides were then washed with distilled water 3 times for 1 minute each, and then a coverslip was mounted using Fluoromount (Electron Microscopy Sciences #17989-70).For NeuN quantification of DAB-stained sections, stitched 20X images covering the hippocampus and cortex of a coronal section were obtained on a Keyence BZ-X800 microscope.

### DiI labeling of dendritic spines

For dendritic protrusion analysis, 400-μm hippocampal-entorhinal cortical slices^1,2^ were prepared as per previously published protocols^1,3^. Slices were cut in a high-sucrose cutting solution containing, in mM, 192 sucrose, 2.5 KCl, 1.25 NaH_2_PO_4_, 26 NaHCO_3_, 12.2 glucose, 3 sodium pyruvate, 5 sodium ascorbate, 2 thiourea, 10 MgSO_4_, and 0.5 CaCl_2_. Slices were allowed to recover for 45 minutes at 37°C and 45 minutes at room temperature before recordings. During recovery, the slices were maintained in artificial cerebrospinal fluid (ACSF) containing, in mM, 115 NaCl, 2.5 KCl, 1.4 NaH_2_PO_4_, 24 NaHCO_3_, 12.5 glucose, 3 sodium pyruvate, 5 sodium ascorbate, 2 thiourea, 1 MgSO_4_, and 2.5 CaCl_2_. For recording, slices were maintained in a standard aCSF solution containing, in mM, 128 NaCl, 2.5 KCl, 1.4 NaH_2_PO_4_, 26.2 NaHCO_3_, 12.2 glucose, 1 MgSO_4_, and 2.5 CaCl_2_.

DiI(1,1′-dioctadecyl-3,3,3′,3′-tetramethylindocarbocyanine perchlorate; catalog no. 468495; Millipore Sigma) crystals were used to fluorescently label neurons as previously described^4^. Slices were placed in 1.5% PFA overnight and then rinsed with PBS three times for 5 min each. DiI crystals were placed precisely over the hippocampal CA1 stratum pyramidale region using thin tungsten wire under a dissection microscope. Slices were covered in parafilm at placed in a humidified chamber for two days at RT. Slices were then further fixed with 4%PFA for 30 min at room temperature. Slices were washed with PBS twice and counterstained with DAPI (Nucblue-R37606, ThermoFisher).

Images for dendrite analyses were acquired using a Leica SP8 Confocal microscope. DiI crystals were visualized using a 561 nm laser at 63X, 1.4 NA, oil immersion, with a Z step size of 0.2um. Slices were obtained from 4 animals per genotype; three slices were analyzed per mouse, with three regions of interest analyzed per slice. Segments of secondary and tertiary dendrites of CA1 pyramidal cells located ∼80um from the soma were selected for analysis. After image acquisition, dendritic spine analysis was performed using the previously published methods^5^. Using ImageJ, nonoverlapping dendrites were analyzed for dendrite number and length.

### EEG Recordings

Recording headcaps were created by attaching 1 ground, 1 reference, cortical leads (0.004 inches, formvar-coated silver, California Fine Wire), and 2 hippocampal leads (0.005 inches bare, 0.008 inches coated, stainless steel wire, AM-Systems) to a microconnector (Omnetics).

For implantation, mice were anesthetized with inhaled isoflurane and given 5mg/kg meloxicam (Pivetal, 21294589) for pain management. The following stereotaxic coordinates (measurements relative to bregma) were used to implant electrodes: bilateral motor cortices: 0.5 mm rostral, 1 mm lateral, and 0.6 mm deep; bilateral barrel field cortices: 0.7 mm caudal, 3 mm lateral, and 0.6 mm deep; bilateral visual cortex: 3.5 mm caudal, 2 mm lateral, and 0.6 mm deep; and bilateral hippocampus (CA1 region): 2.2 mm caudal, 2 mm lateral, and 1.7 mm deep. The cerebellar reference lead was implanted posterior to lambda. The ground wire was wrapped tightly around a 1/8-inch self-tap screw (JI Morris, F000CE125), which was inserted into the skull rostral to the motor cortex leads. To aid in securing the recording cap, a second screw was inserted caudal to lambda. The recording electrodes were secured with dental cement. The mouse was given 5mg/kg meloxicam 24-hours post-operatively and allowed to recover for at least 36 hours before recording.

Electroencephalography data were recorded using the Intan RHS2000 Recording System (Intan Systems, California, USA) at a sampling rate of 2500000 Hz. Data were subsequently downsampled to 2500 Hz. All data analysis was performed in MATLAB (MathWorks, Massachusetts, USA), code is available upon request. Recordings were split into 30-minute time periods.

*Data Preprocessing*. All analyzed data was notch filtered to remove line frequency and passed through a 6^th^ order bandpass Butterworth filter. Detection of poor recording channels was performed using previously reported algorithms^1,6^. In short, the root mean square error (RMSE) and skew of each 30-minute recording period was calculated. Channels were deemed poor for that recording period if the RMSE was less than 30 µV, or greater than 200 µV, or had a skew greater than 0.4. Poor channels were detected with 90% accuracy on a test data set when compared to an expert physician trained to evaluate EEGs using these parameters. Data for each recording period was split into 5 s epochs and normalized for further analysis. Epochs with amplitude z-scores of greater than 3 were considered artifacts and removed for all analyses except for spike detection.

*Spike detection:* Single and multi-lead spikes were detected using previously reported parameters^1,6^. Single spikes were characterized by voltage deflections with z-scores greater than 5 with full widths between ½ the maximum amplitudes of 5-200 ms. Spikes within 10 s of an artifact, a spike outside the 5-200 ms parameters, were ignored. Multi-lead spikes were determined if spikes on two or more leads fired within 100 ms of each other.

*Band and absolute power analysis:* The frequency domain of the normalized data was calculated using the Fast Fourier transform and separating the power spectrum into six major EEG frequency bands (delta 0.1–4.0 Hz, theta 4–8 Hz, alpha 8–13 Hz, beta 13–25 Hz, gamma 25–50 Hz, fast gamma >50 Hz). For band analysis, the absolute power for each 5 s epoch was normalized to the total power for that epoch and averaged across the 30-minute recording period for each frequency band. Absolute power was calculated using the Fast Fourier transform applied over 5 s epochs and averaging for each 30-minute recording period.

---

**Supplementary Table 1.** Amino Acid Standards for LC/MS.

| Unlabeled Calibration Standards | Labeled Internal Standards |
| --- | --- |
| glycine | glycine-D5 |
| L-alanine | DL-alanine-2,3,3,3-D4 |
| L-serine | L-serine-2,3,3-D3 |
| L-proline | L-proline-D7 |
| DL-valine | L-valine-D8 |
| L-leucine | L-leucine-5,5,5-D3 |
| L-isoleucine | L-isoleucine-U-13C6 |
| L-methionine | L-methionine-D3 |
| L-histidine | L-histidine-13C6 hydrochloride monohydrate |
| L-phenylalanine | L-phenylalanine-D8 |
| L-tyrosine | L-tyrosine-Ring-D4 |
| L-asparagine | L-asparagine-13C4 monohydrate |
| L-aspartic acid | L-aspartic acid-2,3,3-D3 |
| L-glutamine | L-glutamine-2,3,3,4,4-D5 |
| L-glutamic acid | DL-glutamic-2,3,3-D3 acid |
| L-ornithine monohydrochloride | L-ornithine-3,3,4,4,5,5-D6 HCl |
| L-arginine | L-arginine-13C6 hydrochloride |
| L-threonine | L-threonine-U-13C4 |
| L-lysine | DL-lysine-4,4,5,5-D4 2HCl |
| L-tryptophan | L-tryptophan-D8 |
| L-citrulline | L-citrulline-4,4,5,5-D4 |

**Supplementary Table 2.**
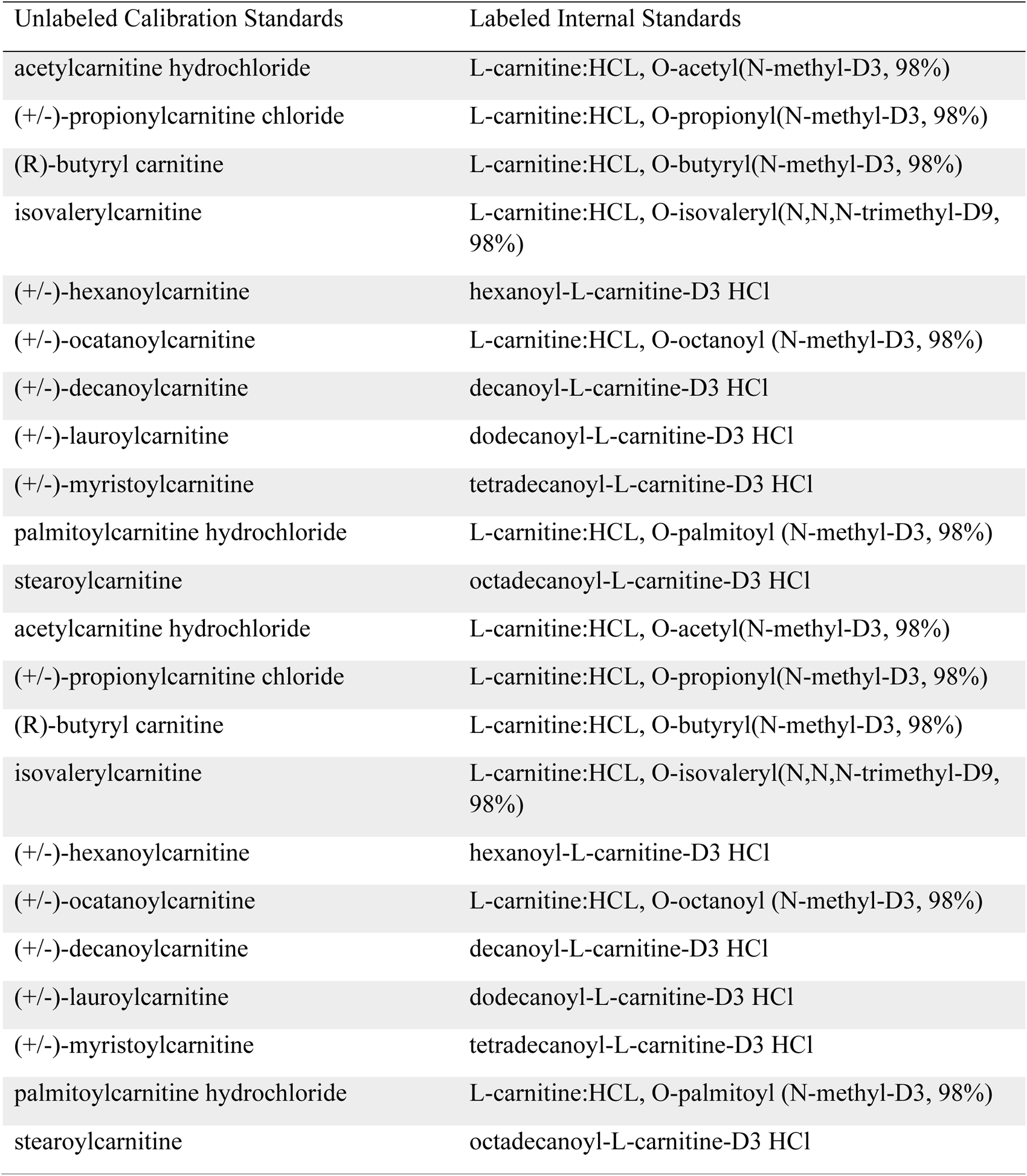
Acylcarnitine Standards for LC/MS.

**Supplementary Figure 1:**
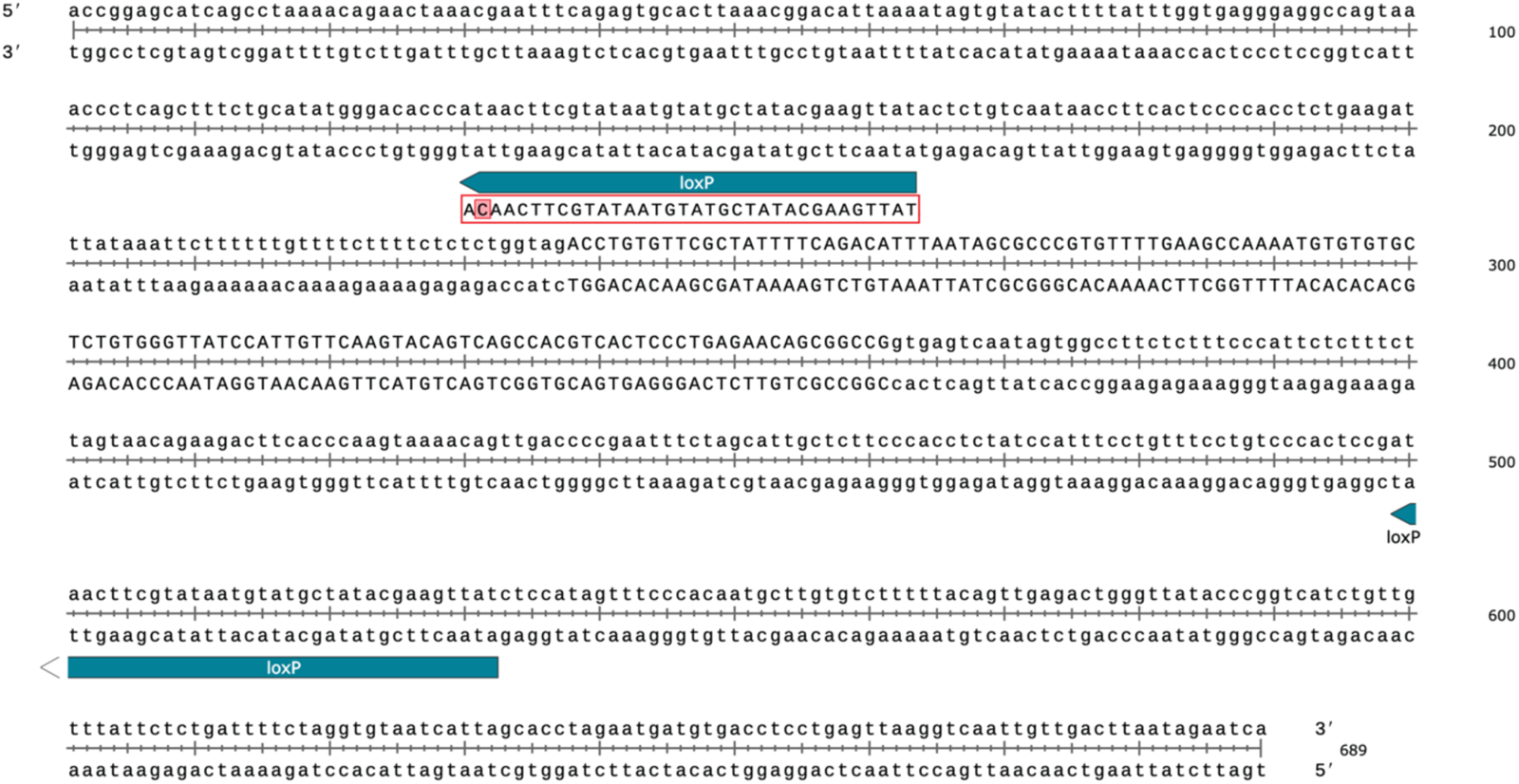
Location of loxP sites introduced into *Dbt* locus. A single missense variant was identified in the proximal loxP sites, as highlighted in red. This variant did not prevent successful cre-mediated recombination in *Dbt*^Flox/Flox^ mice.

**Supplementary Figure 2:**
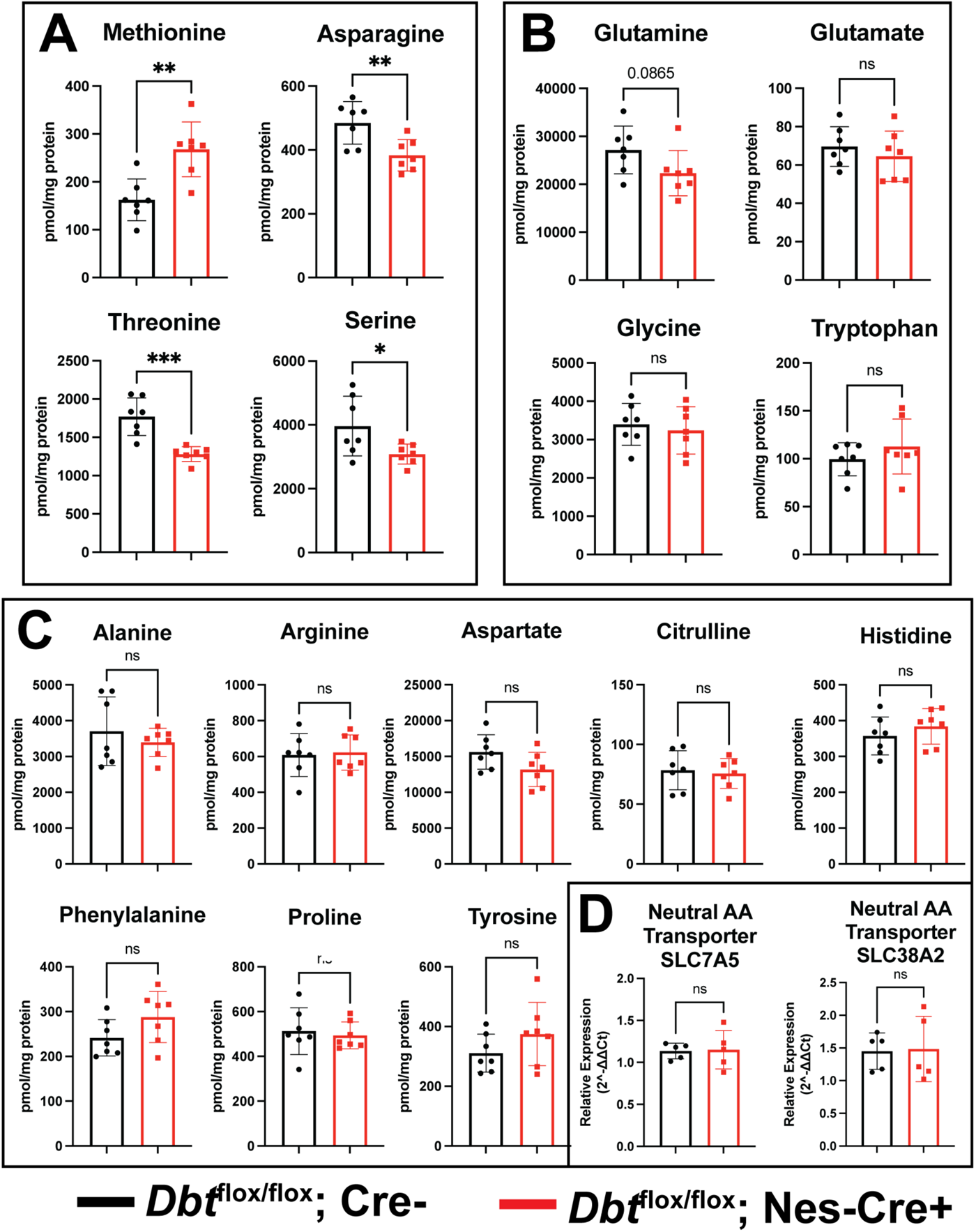
Additional amino acid and transporter levels in p30 cortex from brain KO MSUD mice. **(A)** A small number of non-BCAA amino acids showed significant changes in concentration in brain KO MSUD mice (red) as compared to controls (black). **(B)** Key CNS amino acids did not significantly differ between brain KO MSUD and control mice, neither did **(C)** additional AAs evaluated. **(D)** Expression of neutral AA transporters (as measured at the transcript level) was also unchanged. Data shown are mean +/-standard error, and groups were compared via student’s t-test with *p<0.05, **p<0.01, ***p<0.001, ****p<0.0001.

**Supplementary Figure 3:**
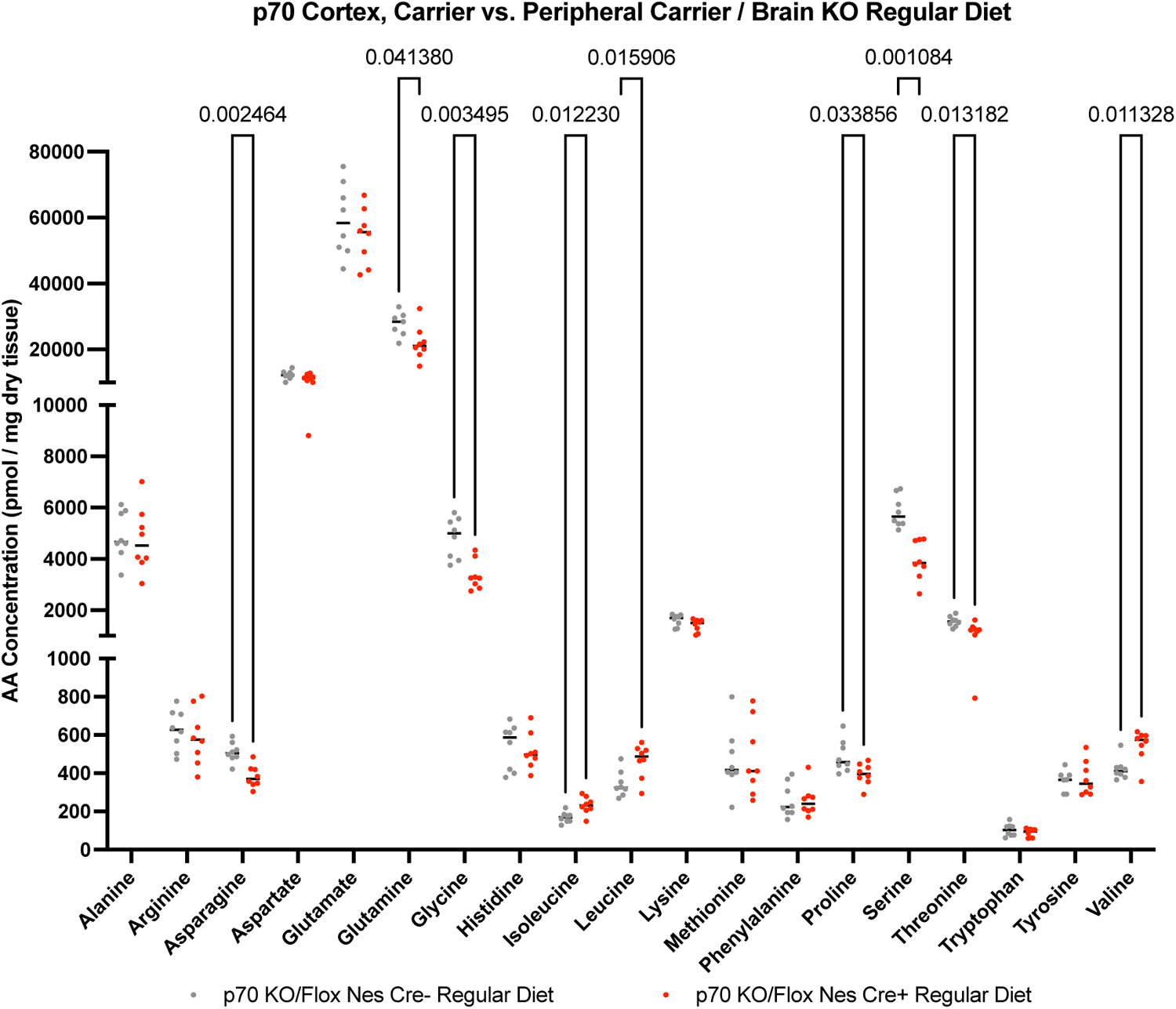
Amino acid levels in cortex of p70 peripheral carrier, brain KO MSUD mouse model on a regular protein diet. P70 peripheral carrier, brain KO mice were investigated as a model of partially treated MSUD (such as occurs after liver transplantation). Amino acid levels in cortex of peripheral carrier, brain KO MSUD mice on a regular (20%) protein diet were compared to carrier control mice on a regular diet. Between group differences were analyzed by multiple t-tests with multiple comparison analysis using the two-stage step-up method of Benjamini, Krieger and Yekutieli with a desired q of 5.00%, with significant q-values shown. N=8 animals per genotype group.

**Supplementary Figure 4:**
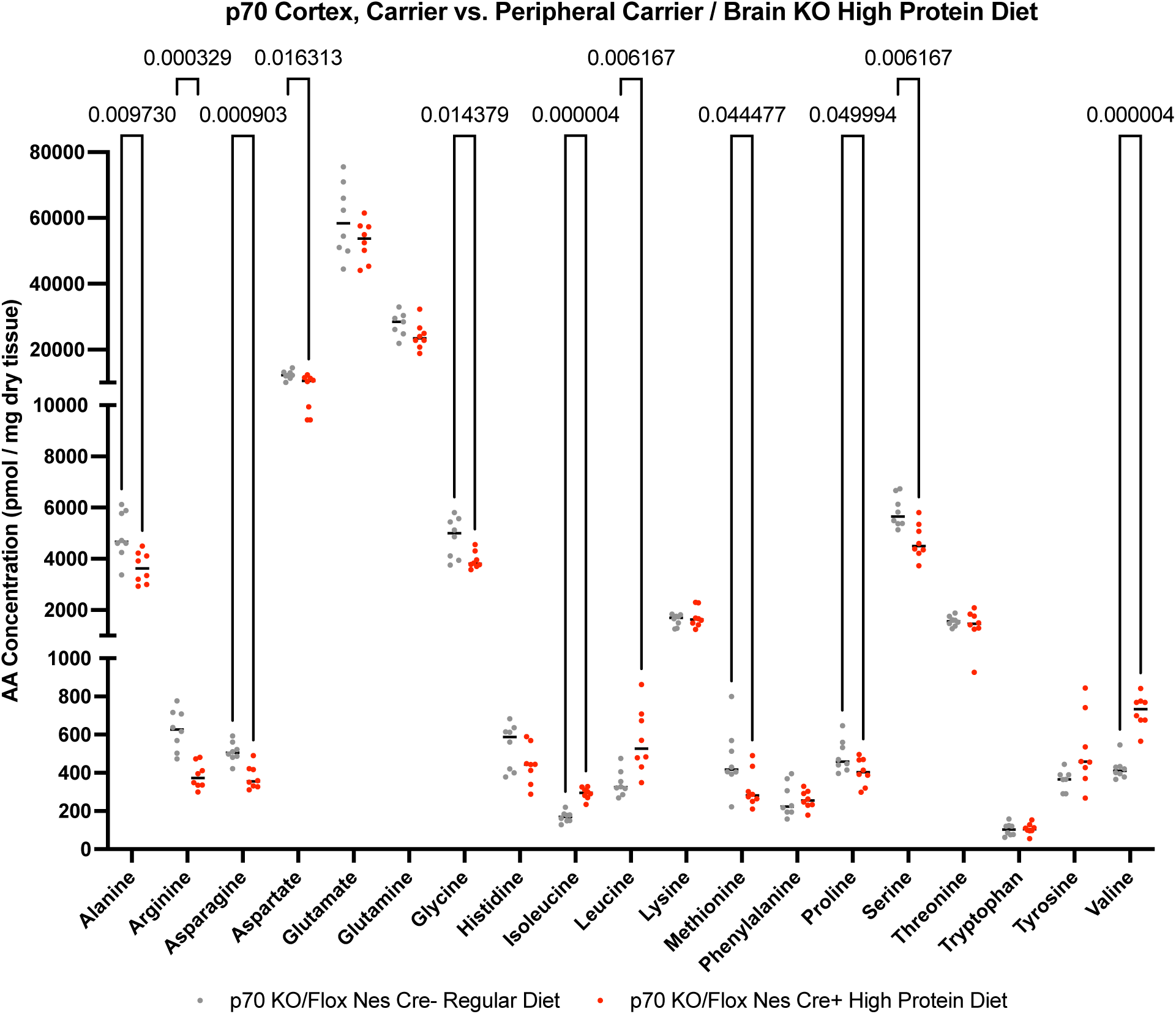
Amino acid levels in cortex of p70 peripheral carrier, brain KO MSUD mouse model on a high protein diet. P70 peripheral carrier, brain KO mice were investigated as a model of partially treated MSUD (such as occurs after liver transplantation). Amino acid levels in cortex of peripheral carrier, brain KO MSUD mice on a high (20%) protein diet were compared to carrier control mice on a regular diet. Between group differences were analyzed by multiple t-tests with multiple comparison analysis using the two-stage step-up method of Benjamini, Krieger and Yekutieli with a desired q of 5.00%, with significant q-values shown. N=8 animals per genotype group.

**Supplementary Figure 5:**
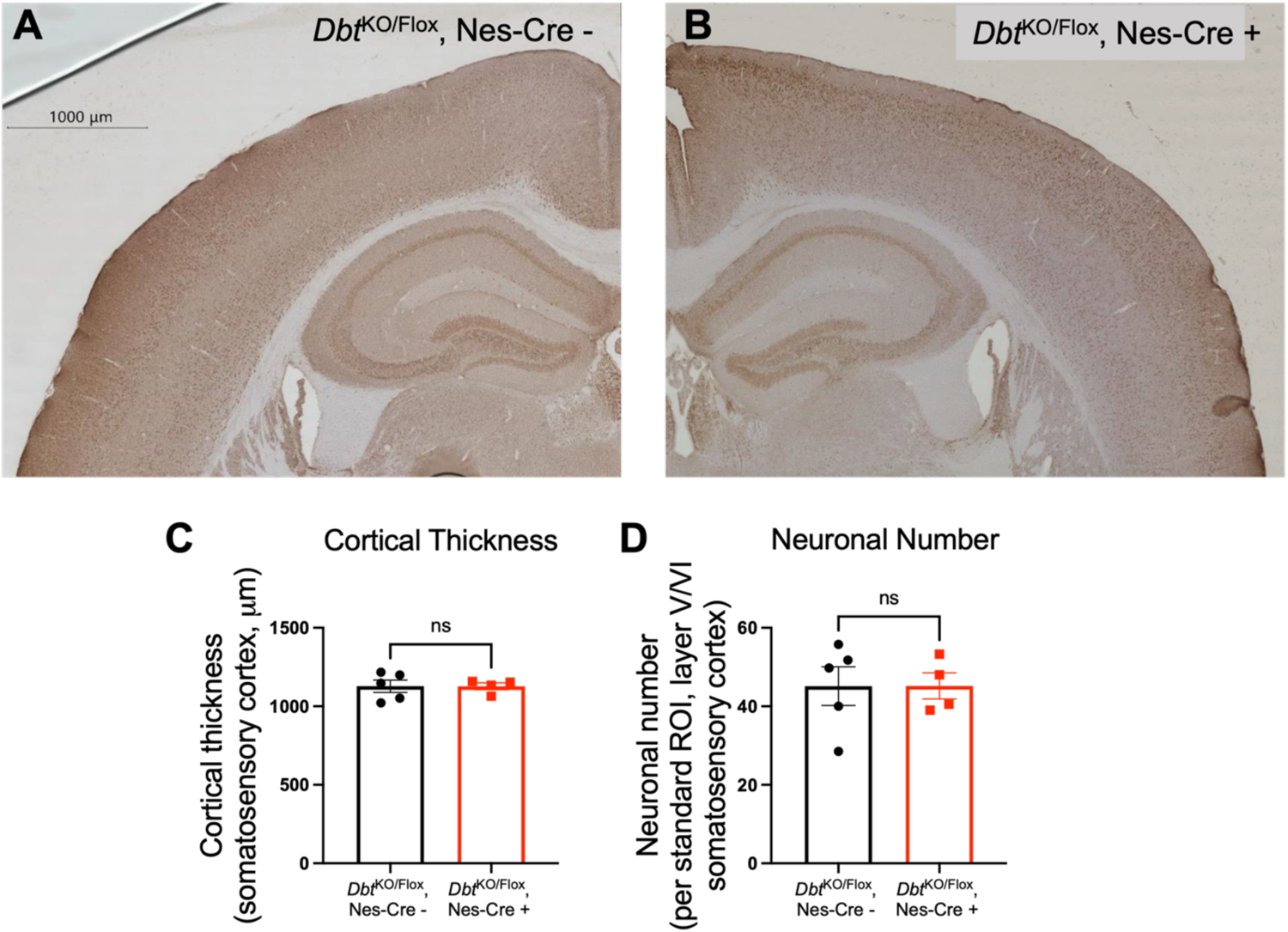
Peripheral carrier, brain KO MSUD mice have normal cortical thickness and neuronal number. Immunohistochemistry to detect NeuN, a pan-neuronal marker, was completed in **(A)** peripheral carrier control (*Dbt*^KO/Flox^, Cre -) and **(B)** peripheral carrier, brain KO MSUD mice. **(C)** Cortical thickness was measured and averages at four standard locations for each animal. No differences were detected between genotypes. **(D)** Neuronal number was calculated from three consistently paced and sized regions of interest in layer 5/6 of the somatosensory cortex. No differences were detected by genotype. Data shown as mean +/-standard error. Groups were compared using student’s t-test.

**Supplementary Figure 6:**
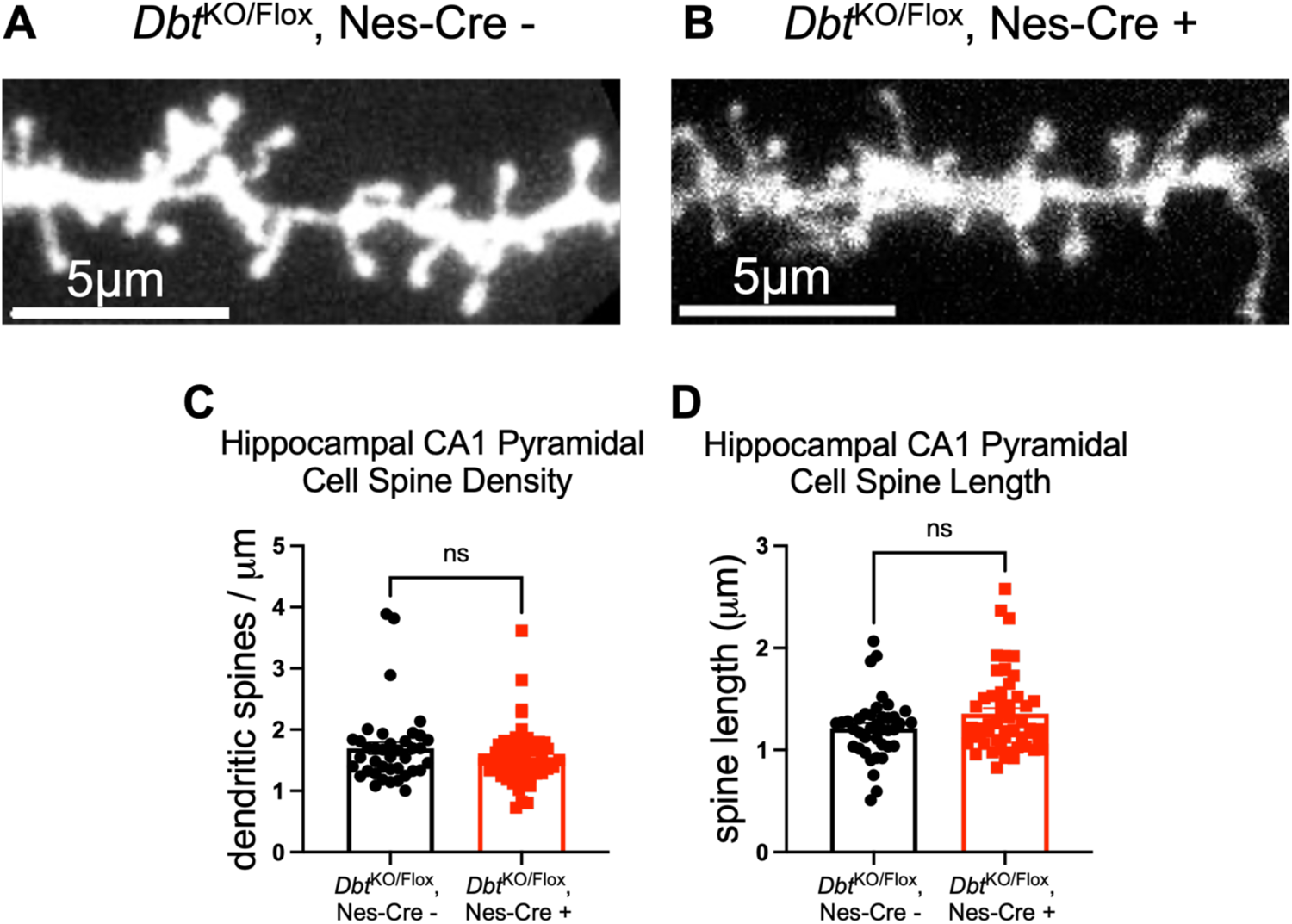
Peripheral carrier, brain KO MSUD mice have normal dendritic spine density and length in the CA1 region of the hippocampus. DiI-labeling of dendritic protrusions in the CA1 region of the hippocampus of **(A)** peripheral carrier control (*Dbt*^KO/Flox^, Cre -) and **(B)** peripheral carrier, brain KO MSUD mice. **(C)** There were no differences in spine density or **(D)** spine length. Data shown as mean +/-standard error. Groups were compared using student’s t-test. N= a minimum of 35 dendrites from 4 mice per genotype group.

**Supplementary Figure 7:**
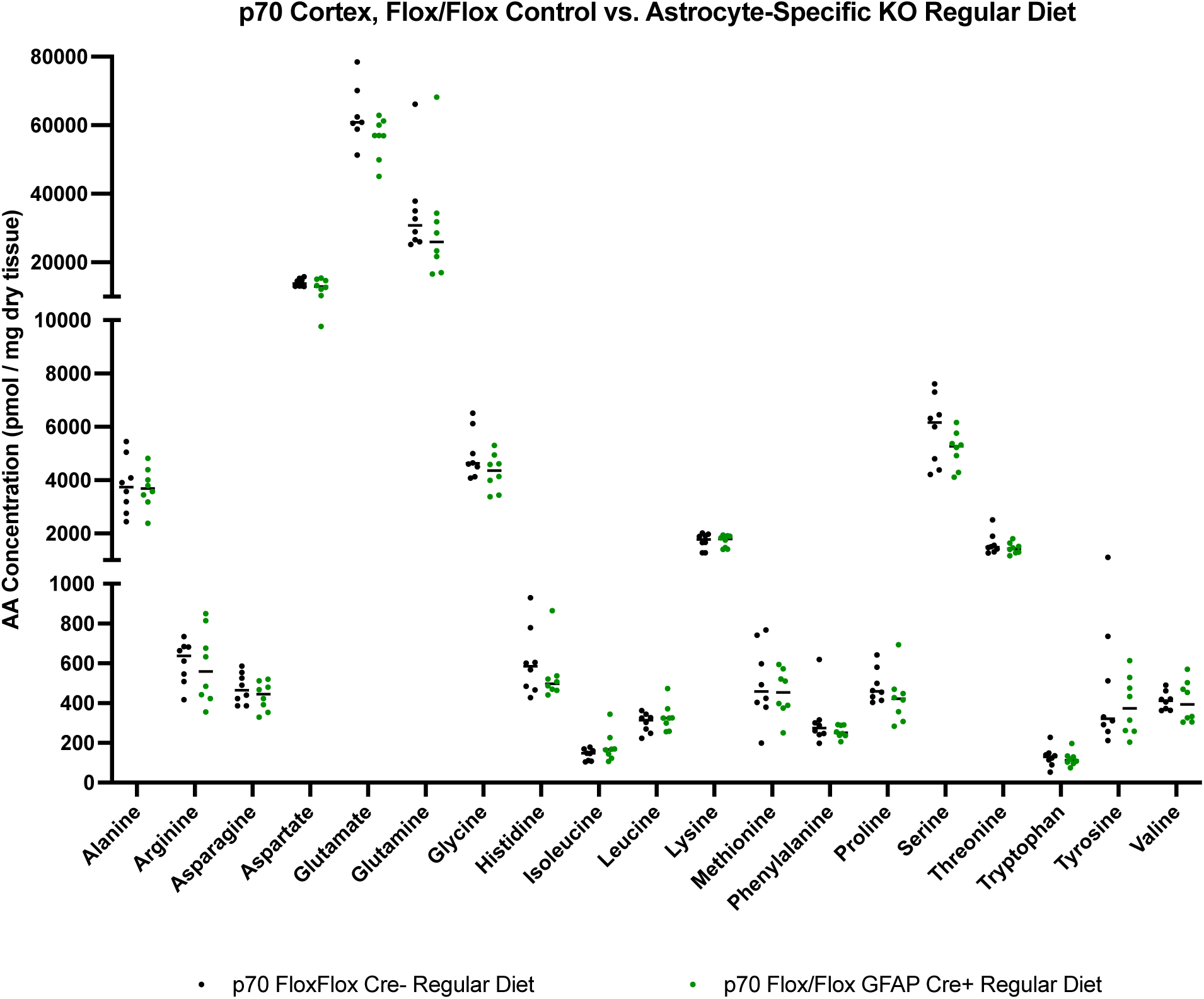
Amino acid levels in cortex of p70 astrocyte KO MSUD mouse model on a regular protein diet. Amino acid levels in cortex of astrocyte-specific KO MSUD mice on a regular (20%) protein diet were compared to *Dbt*^Flox/Flox^, Cre-control mice on a regular diet. Between group differences were analyzed by multiple t-tests with multiple comparison analysis using the two-stage step-up method of Benjamini, Krieger and Yekutieli with a desired q of 5.00%, with significant q-values shown. N=8 animals per genotype group.

**Supplementary Figure 8:**
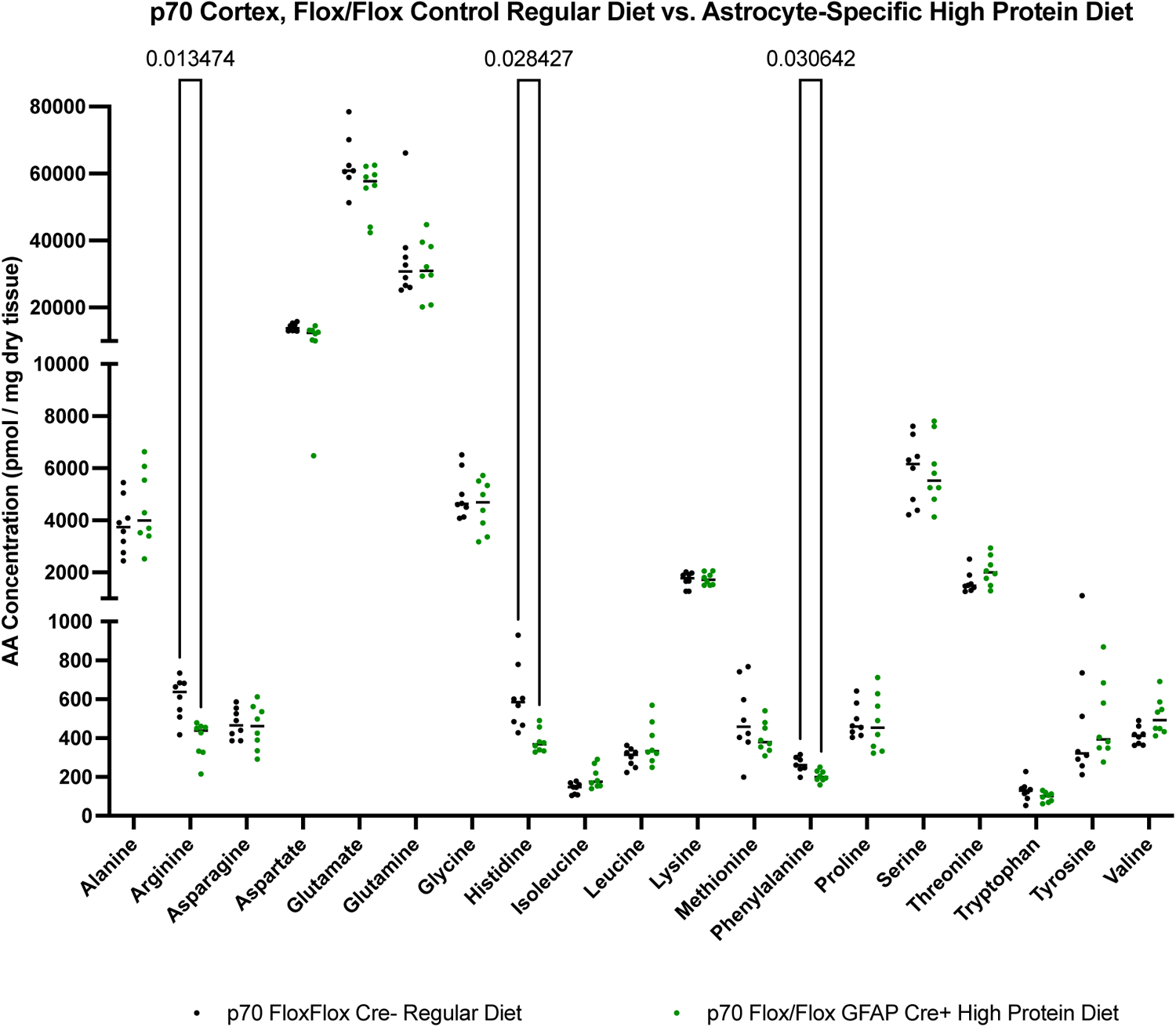
Amino acid levels in cortex of p70 astrocyte KO MSUD mouse model on a high protein diet. Amino acid levels in cortex of astrocyte-specific KO MSUD mice on a high (40%) protein diet were compared to *Dbt*^Flox/Flox^, Cre-control mice on a regular diet. Between group differences were analyzed by multiple t-tests with multiple comparison analysis using the two-stage step-up method of Benjamini, Krieger and Yekutieli with a desired q of 5.00%, with significant q-values shown. N=8 animals per genotype group.

**Supplementary Figure 9:**
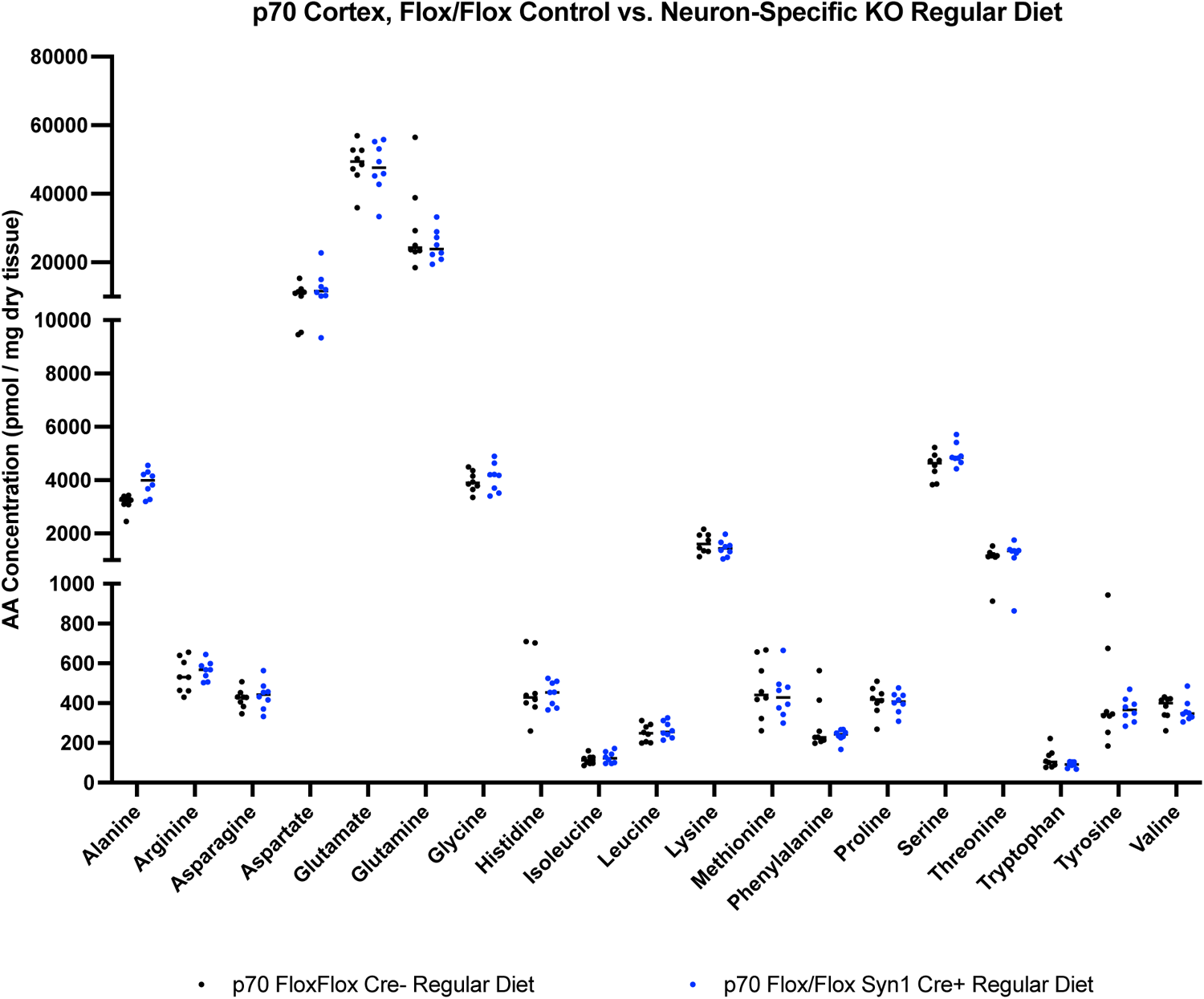
Amino acid levels in cortex of p70 neuron KO MSUD mouse model on a regular protein diet. Amino acid levels in cortex of neuron-specific KO MSUD mice on a regular (20%) protein diet were compared to *Dbt*^Flox/Flox^, Cre-control mice on a regular diet. Between group differences were analyzed by multiple t-tests with multiple comparison analysis using the two-stage step-up method of Benjamini, Krieger and Yekutieli with a desired q of 5.00%, with significant q-values shown. N=8 animals per genotype group.

**Supplementary Figure 10:**
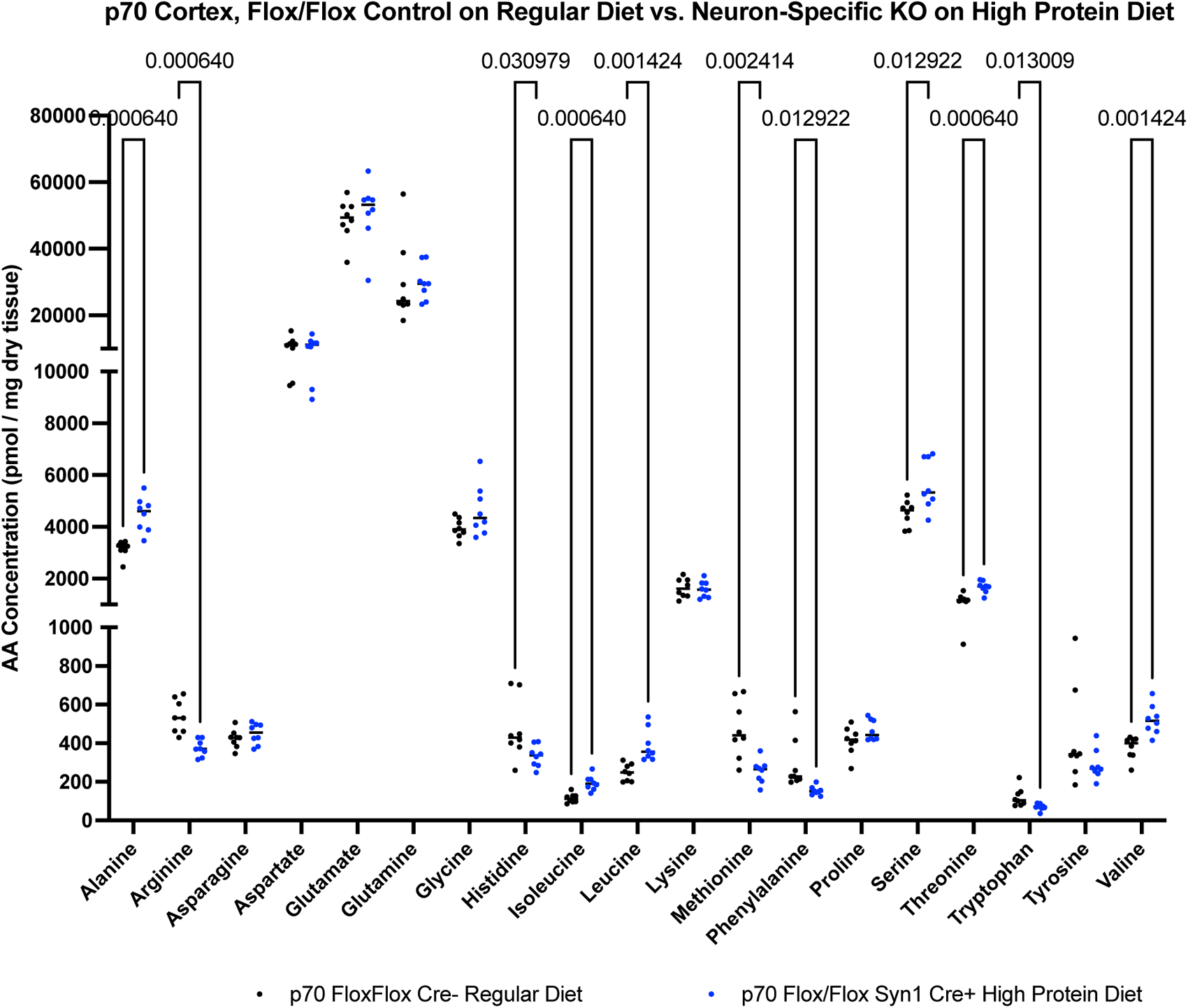
Amino acid levels in cortex of p70 neuron KO MSUD mouse model on a high protein diet. Amino acid levels in cortex of neuron-specific KO MSUD mice on a high (40%) protein diet were compared to *Dbt*^Flox/Flox^, Cre-control mice on a regular diet. Between group differences were analyzed by multiple t-tests with multiple comparison analysis using the two-stage step-up method of Benjamini, Krieger and Yekutieli with a desired q of 5.00%, with significant q-values shown. N=8 animals per genotype group.

**Supplementary Figure 11:**
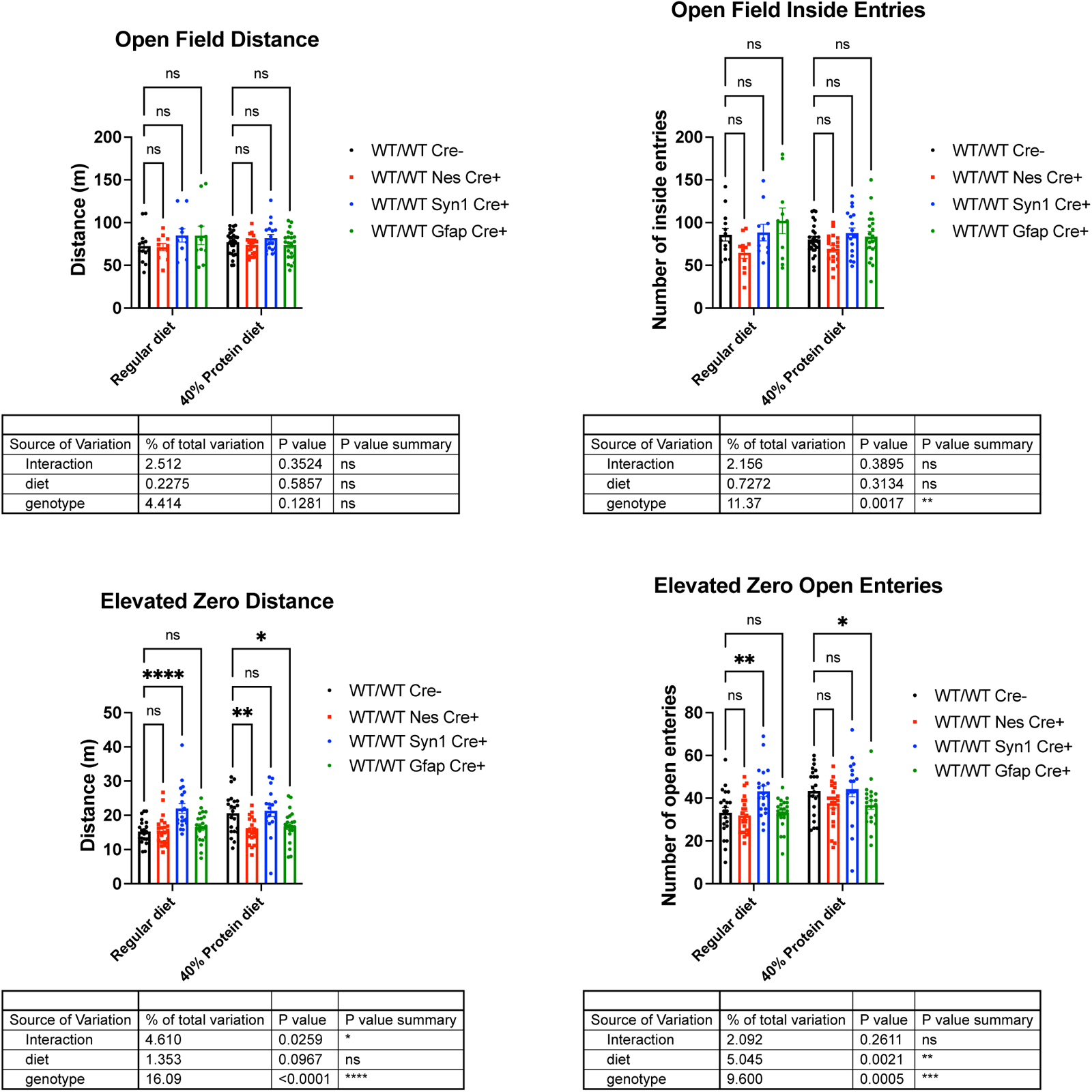
Cre driver alleles affect behavioral phenotypes on elevated zero testing, but have minimal impact on open field testing. (Top) Wildtype mice carrying the Cre driver alleles employed in this study had minimal effect on distance traveled on open field testing or entries into the inside zone. **(Bottom)** Conversely, the Cre driver alleles influenced both distance traveled and entries into the open zone of an elevated zero maze. N= a minimum of 17 animals per genotype group. Groups compared via 2-way ANOVA followed by Sidak’s multiple comparison testing with *p<0.05, **p<0.01, ***p<0.001, ****p<0.0001.

